# Phospho-regulation accommodates Type III secretion and assembly of a tether of ER-*Chlamydia* inclusion membrane contact sites

**DOI:** 10.1101/2021.10.19.465067

**Authors:** Rebecca L. Murray, Rachel J. Ende, Samantha K. D’Spain, Isabelle Derré

**Affiliations:** Department of Microbiology, Immunology, and Cancer Biology, University of Virginia School of Medicine, Charlottesville, Virginia, USA

## Abstract

Membrane contact sites (MCS) are crucial for non-vesicular trafficking-based inter-organelle communication. ER-organelle tethering occurs in part through the interaction of the ER resident protein VAP with FFAT-motif containing proteins. FFAT motifs are characterized by a seven amino acidic core surrounded by acid tracks. We have previously shown that the human intracellular bacterial pathogen *Chlamydia trachomatis* establishes MCS between its vacuole (the inclusion) and the ER through expression of a bacterial tether, IncV, displaying molecular mimicry of eukaryotic FFAT motif cores. Here, we show that multiple layers of host cell kinase-mediated phosphorylation events govern the assembly of the IncV-VAP tethering complex. CK2-mediated phosphorylation of a C-terminal region of IncV enables IncV hyperphosphorylation of a phospho-FFAT motif core and serine-rich tracts immediately upstream of IncV FFAT motif cores. Phosphorylatable serine tracts, rather than genetically-encoded acidic tracts, accommodate Type III-mediated translocation of IncV to the inclusion membrane, while achieving full mimicry of FFAT motifs. Thus, regulatory components and post-translational modifications are integral to MCS biology, and intracellular pathogens such as *C. trachomatis* have evolved complex molecular mimicry of these eukaryotic features.

## Introduction

In naïve cells, membrane contact sites (MCS) are points of contact between the membrane of two adjacent organelles (10-30 nm apart). They provide physical platforms for the non-vesicular transfer of lipids and ions, and cell signaling events important for inter-organelle communication and organelle positioning and dynamics (Prinz et al., 2020). Since their discovery and implication in cell homeostasis, MCS dysfunction has been linked to several human diseases (Area-Gomez et al., 2012; Castro et al., 2018; Stoica et al., 2014). At the molecular level, depending on the contacting organelles [endoplasmic reticulum (ER)-Golgi, ER-mitochondria, ER-plasma membrane (PM), etc…], each MCS is enriched in specific proteinaceous factors that contribute to the specialized biological function of a given MCS (Prinz et al., 2020). By bridging the membrane of apposed organelles, either via protein-protein or protein-lipid interactions, MCS components also form tethering complexes that increase the affinity of one organelle to another and thereby keep their membranes in close proximity (Eisenberg-Bord et al., 2016; Prinz et al., 2020; Scorrano et al., 2019). Although the overall molecular composition of each MCS is different, one integral ER protein, the vesicle-associated membrane protein (VAMP)-associated protein (VAP) (Murphy & Levine, 2016), engages in tethering complexes at several MCS. This is accomplished by interaction of the cytosolic major sperm protein (MSP) domain of VAP with proteins containing two phenylalanine (FF) in an acidic tract (FFAT) motifs (Loewen et al., 2003; Murphy & Levine, 2016). FFAT motif containing proteins include soluble proteins, such as lipid transfer proteins that contain an additional domain for targeting to the opposing membrane, and transmembrane proteins anchored to the contacting organelle (James & Kehlenbach, 2021). The molecular determinants driving the VAP-FFAT interaction have been investigated at the cellular and structural level. A consensus of the FFAT motif core was first defined as seven amino acids, E^1^F^2^F^3^D^4^A^5^x^6^E^7^; however, the core motif of many identified VAP interacting proteins deviates from this canonical sequence (James & Kehlenbach, 2021; Loewen et al., 2003). In addition to the core, acidic residues surrounding the core motif are proposed to facilitate the VAP-FFAT interaction through electrostatic interactions (Furuita et al., 2010).

In addition to their critical role in inter-organelle communication, MCS are exploited by intracellular pathogens for replication (Derré, 2017; Ishikawa-Sasaki et al., 2018; Justis et al., 2017; McCune et al., 2017). One example is the obligate intracellular bacterium *Chlamydia trachomatis,* the causative agent of the most commonly reported bacterial sexually transmitted infection. Upon invasion of the genital epithelium, *C. trachomatis* replicates within a membrane-bound vacuole called the inclusion (Gitsels et al., 2019). Maturation of the inclusion relies on *Chlamydia* effector proteins that are translocated across the inclusion membrane *via* a bacterially encoded Type III secretion system (Lara-Tejero & Galan, 2019). A subset of *Chlamydia* Type III effector proteins, known as the inclusion membrane proteins (Inc), are inserted into the inclusion membrane and are therefore strategically positioned to mediate inclusion interactions with host cell organelles (Bugalhao & Mota, 2019; Dehoux et al., 2011; Lutter et al., 2012; Moore & Ouellette, 2014). These interactions include points of contact between the ER and the inclusion membrane, without membrane fusion (Derre et al., 2011; Dumoux et al., 2012), which are referred to as ER-Inclusion MCS based on their similarities to MCS between cellular organelles (Agaisse & Derre, 2015; Derre et al., 2011).

Characterization of the protein composition of ER-Inclusion MCS led to the identification the Inc protein IncV, which constitutes a structural component that tethers the ER membrane to the inclusion membrane through interaction with VAP (Stanhope et al., 2017). The IncV-VAP interaction relies on the presence of two FFAT motifs in the C-terminal cytosolic tail of IncV. The core sequence of one of the motifs (_286_E^1^Y^2^M^3^D^4^A^5^L^6^E_7292_) is similar to the canonical sequence, whereas a second motif (_262_S^1^F^2^H^3^T^4^P^5^P^6^N_7268_) deviates significantly and was originally defined as a non-canonical FFAT (Stanhope et al., 2017). Similar to eukaryotic FFAT, the residue in position 2 in each motif (Y_287_ and F_263_, respectively) are essential for the IncV-VAP interaction during infection. However, it remains unclear whether additional determinants promote the assembly of this bacterial tether.

Here, we show that multiple layers of host cell kinase-mediated phosphorylation govern the assembly of the IncV-VAP tethering complex. IncV phosphorylation supports the IncV-VAP interaction through FFAT motifs displaying core domains immediately downstream of phosphorylation-mediated acidic tracts. Since the substitution for genetically encoded acidic tracts interfered with IncV translocation, we propose that *Chlamydia* evolved a post-translocation phosphorylation strategy in order to accommodate proper secretion via the Type III secretion system, while achieving full mimicry of eukaryotic FFAT motifs.

## Results

### A phospho-FFAT motif in IncV contributes to the IncV-VAP interaction

We have previously shown that IncV displays one non-canonical (_262_S^1^F^2^H^3^T^4^P^5^P^6^N_7268_) and one canonical FFAT motif (_286_E^1^Y^2^M^3^D^4^A^5^L^6^E_7292_) (Fig. 1A) (Stanhope et al., 2017). In agreement with position 2 of a FFAT motif being a phenylalanine or a tyrosine residue critical for VAP-FFAT interactions (Kawano et al., 2006; Loewen et al., 2003), we had shown that alanine substitution of residue in position 2 of each motif, individually (IncV_F263A_ or IncV_Y287A_) and in combination (IncV_F263A/Y287A_), led to a partial and full reduction of the IncV-VAP interaction, respectively, indicating that both FFAT motifs cooperate for VAP binding (Stanhope et al., 2017). Recently, Di Mattia et al., identified a new class of FFAT motifs referred to as phospho-FFAT motifs in which the acidic residue in position 4 is replaced by a phosphorylatable residue, such as serine or threonine (Di Mattia et al., 2020). The presence of a phosphorylatable threonine residue in position 4 of the non-canonical FFAT motif of IncV (T_265_) (Fig. 1A) suggests that, as proposed by Di Mattia et al., the non-canonical FFAT of IncV is a phospho-FFAT motif. To test this hypothesis, T_265_ was substituted for an alanine residue either individually (IncV_T265A_), or in combination with alanine mutation of the tyrosine residue in position two of the canonical FFAT (IncV_T265A/Y287A_). HeLa cells expressing YFP-VAP were infected with a previously characterized *incV* mutant strain of *C. trachomatis* (Stanhope et al., 2017; Weber et al., 2017), expressing IncV_T265A_- or IncV_T265A/Y287A_- 3xFLAG under the control of the anhydrotetracycline (aTc)-inducible promoter. Cells infected with *incV* mutant strains expressing IncV_WT_-, IncV_Y287A_-, and IncV_F263A/Y287A_-3xFLAG were included as controls. The cells were fixed at 24 h post infection, immunostained with anti-FLAG antibody, and analyzed by confocal immunofluorescence microscopy (Fig. 1B). All IncV constructs were equally localized to the inclusion membrane (Fig. S1A). As previously observed (Stanhope et al., 2017), IncV_WT_ exhibited a strong association of YFP-VAP with the inclusion membrane, while IncV_Y287A_ and IncV_F263A/Y287A_ exhibited a significant partial and full loss of inclusion associated YFP-VAP, respectively (Fig. 1B-C). Similarly, IncV _T265A_ and IncV_T265A/Y287A_ exhibited partial and complete loss of VAP association with the inclusion, respectively (Fig. 1B-C) indicating that mutation of residues at position 4 (T_265_) of IncV non-canonical FFAT motif is critical to mediate the VAP-FFAT interaction. Altogether, these results experimentally validate that the non-canonical FFAT motif of IncV is a phospho-FFAT motif and identify T_265_ as a core residue mediating the IncV-VAP interaction.

**Figure 1:**
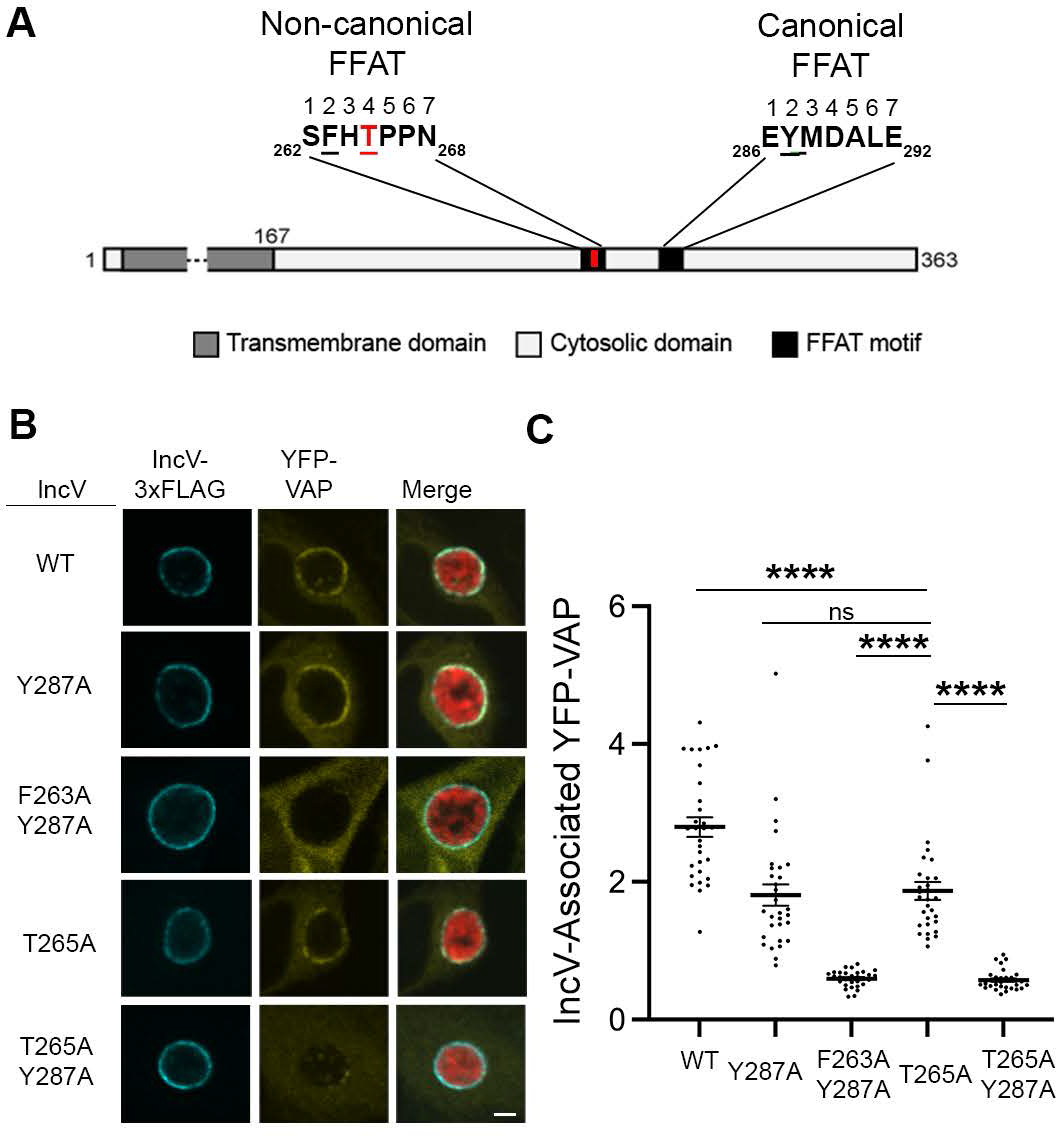
A phospho-FFAT motif in IncV contributes to the IncV-VAP interaction. (A) Schematic depicting the IncV protein. The transmembrane domain, the cytosolic domain, and the non-canonical and the canonical FFAT motif cores are indicated in dark grey, light grey and black, respectively. The amino acid sequence of the FFAT motif cores is shown. Numbers 1-7 indicate the amino acid position within the FFAT motif cores, other numbers indicate the amino acid position within the IncV protein sequence. Residues at position 2 of the FFAT motif cores are in black and underlined. Threonine 265 at position 4 of the non-canonical FFAT is in red and underlined. (B) Single plane confocal images of HeLa cells expressing YFP-VAP (yellow), infected with a *C. trachomatis incV* mutant expressing mCherry constitutively (red) and IncV_WT_- 3xFLAG (WT), IncV_Y287A_-3xFLAG (Y287A), IncV_F263A/Y287A_ (F263A/Y287A), IncV_T265A_-3xFLAG (T265A), or IncV_T265A/Y287A_-3xFLAG (T265A/Y287A) (blue) under the control of an aTc inducible promoter. The merge is shown on the right. Scale bar is 5μm. (C) Quantification of the mean intensity of YFP-VAP within an object generated from the IncV-3xFLAG signal and normalized to the mean intensity of YFP-VAP in the ER. Each dot represents one inclusion. Data show the mean and SEM of a representative experiment. One-way ANOVA and Tukey’s post hoc test was performed. **** *P* <0.0001.

### IncV is modified by phosphorylation

The presence of a phospho-FFAT in IncV led us to investigate the phosphorylation status of IncV. When subjected to anti-FLAG western blot analysis, lysates of HEK293 eukaryotic cells infected with wild type *C. trachomatis* expressing IncV-3xFLAG displayed a doublet consisting of a 50kDa and 60 kDa band (Fig. 2A, middle lane, 293 + *Ct*). By contrast, IncV-3xFLAG ectopically expressed in HEK293 cells had an apparent molecular weight that was shifted toward the 60k Da band of the doublet (Fig. 2A, left lane, 293), while IncV-3xFLAG expressed in *E. coli* had an apparent molecular weight equivalent to the 50 kDa band of the doublet (Fig. 2A, right lane, *Ec*). This result led us to hypothesize that IncV is post-translationally modified by a host factor.

**Figure 2:**
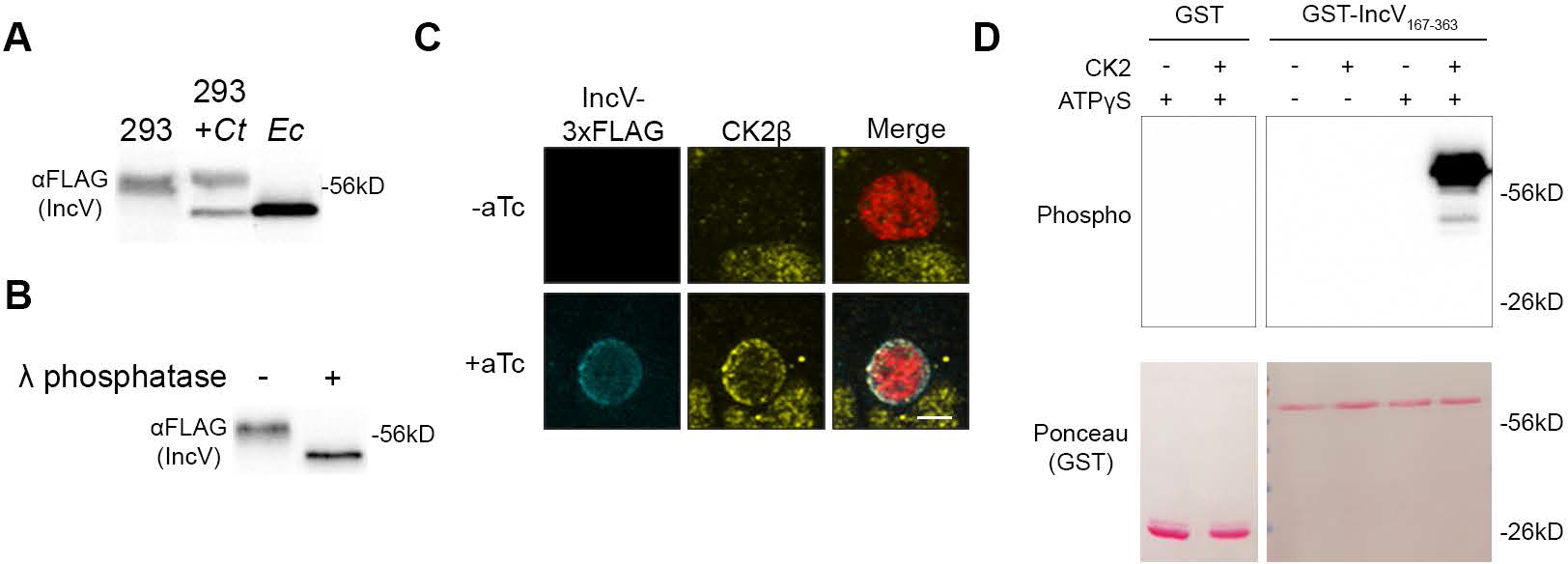
CK2 localize to the inclusion and phosphorylates IncV. (A) Western blot of IncV- 3xFLAG from lysates of HEK293 cells expressing IncV-3xFLAG (293), HEK293 cells infected with *C. trachomatis* expressing IncV-3xFLAG (293 + *Ct*), or *E. coli* expressing IncV-3xFLAG (*Ec*). (B) Western blot of IncV-3xFLAG purified from lysates of HEK293 cells infected with *C. trachomatis* expressing IncV-3xFLAG and treated with lambda (λ) phosphatase (+) or phosphatase buffer alone (-). (C) 3-dimensional reconstruction of confocal images of HeLa cells infected with *C. trachomatis* expressing mCherry constitutively (red) and IncV-3xFLAG (blue) under the control of an anhydrotetracycline (aTc)-inducible promoter in the absence (-aTc) or presence (+aTc) of aTc and stained to detect endogenous CK2β (Yellow). The merge is shown on the right. Scale bar is 5μm. (D) *In vitro* kinase assay using GST or GST-IncV_167-363_ purified from *E. coli* as a substrate in the presence (+) or absence (-) of recombinant CK2 and in the presence (+) or absence (-) of ATPγS. The top panel shows phosphorylated proteins detected with anti-Thiophosphate antibodies and the bottom panel is the same membrane stained with Ponceau S to detect total proteins.

To determine if phosphorylation could account for the increase in the apparent molecular weight of IncV, we performed a phosphatase assay. IncV-3xFLAG was immunoprecipitated, using anti-FLAG-conjugated Sepharose beads, from lysates of HEK293 cells infected with *C. trachomatis* expressing IncV-3xFLAG. Following the release of IncV-3xFLAG from the beads by FLAG peptide competition, the eluate was treated with lambda (λ) phosphatase or phosphatase buffer alone, and subsequently subjected to anti-FLAG western blot analysis (Fig. 2B). In the absence of λ phosphatase, the apparent molecular weight of IncV-3xFLAG was approximately 60 kDa (Fig. 2B, left lane). Upon phosphatase treatment, we observed a decrease in the apparent molecular weight of IncV-3xFLAG to approximately 50 kDa, similar to what was observed when IncV- 3xFLAG was expressed in *E. coli* (Fig. 2B, right lane). Altogether, these results demonstrate that IncV is phosphorylated by a host cell kinase.

### The host kinase CK2 phosphorylates IncV

We next focused on identifying the host cell kinase(s) responsible for phosphorylating IncV. All three subunits of Protein Kinase CK2 were identified as potential interacting partners of IncV in an Inc-human interactome (Mirrashidi et al., 2015). To determine if CK2 associated with IncV at ER-Inclusion MCS, HeLa cells transfected with YFP-CK2α or YFP-CK2β constructs and infected with *C. trachomatis* wild type expressing mCherry under a constitutive promoter and IncV- 3xFLAG under the aTc-inducible promoter were analyzed by confocal immunofluorescence microscopy (Fig. S2). In the absence of IncV-3xFLAG expression, YFP-CK2α and YFP-CK2β were undetectable at the inclusion (Fig. S2A and S2B, -aTc). However, upon expression of IncV- 3xFLAG, YFP-CK2α and YFP-CK2β were recruited to the inclusion membrane and colocalized with IncV (Fig. S2A and S2B, +aTc). To confirm that this phenotype was not the result of overexpression of the CK2 subunits, we used antibodies that recognized the endogenous CK2β subunit and showed that endogenous CK2β colocalized with IncV at the inclusion, when IncV- 3xFLAG expression was induced (Fig. 2C). Altogether, these results demonstrate that CK2 is a novel component of ER-Inclusion MCS that is recruited to the inclusion in an IncV-dependent manner.

Having established that IncV is phosphorylated and that CK2 localizes to ER-Inclusion MCS in an IncV-dependent manner, we next tested if CK2 phosphorylates IncV. We performed an *in vitro* kinase assay using recombinant CK2 and the cytosolic domain of IncV (amino acids 167-363 of IncV) fused to GST (GST-IncV_167-363_) or GST alone, purified from *E. coli*. To detect phosphorylation, we used ATPγS, which can be utilized by kinases to thiophosphorylate a substrate, followed by an alkylation reaction of the thiol group to generate an epitope that is detected using an antibody that recognizes thiophosphate esters (Allen et al., 2007). When GST alone was provided as a substrate, there was no detectable phosphorylation, regardless of the presence of CK2 and ATPγS (Fig. 2D, lanes 1 and 2). A similar result was observed with GST- IncV_167-363_ in the absence of CK2 and/or ATPγS (Fig. 2D, lanes 3 - 5). However, in the presence of both ATPγS and CK2, GST-IncV_167-363_ was phosphorylated (Fig. 2D, lane 6). Altogether, these results demonstrate that CK2 directly phosphorylates IncV *in vitro*.

### Phosphorylation of IncV is necessary and sufficient to promote the IncV-VAP interaction *in vitro*

We have previously reported an IncV-VAP interaction *in vitro* upon incubation of IncV_167-363_ with the cytosolic MSP domain of VAP (GST-VAP_MSP_) purified from *E. coli* (Stanhope et al., 2017). However, this interaction was only detected when IncV_167-363_ was produced in eukaryotic cells, which, based on the above results, led us to hypothesize that IncV phosphorylation is required for the IncV-VAP interaction. We assessed the role of phosphorylation in the IncV-VAP interaction by performing lambda (λ) phosphatase dephosphorylation of IncV coupled with a GST-VAP_MSP_ pull-down assay (Fig. 3A). IncV-3xFLAG was immunoprecipitated from lysates of HEK293 cells using anti-FLAG-conjugated Sepharose beads, released from the beads using FLAG peptide competition, and treated with λ phosphatase or buffer alone. Treated and untreated IncV-3xFLAG samples were then incubated with GST-VAP_MSP_ or GST alone bound to glutathione Sepharose beads. The protein-bound beads were subjected to western blot analysis using an anti-FLAG antibody (Fig. 3B). Untreated IncV-3xFLAG was pulled down by GST-VAP_MSP_ but not by GST alone, demonstrating a specific interaction between IncV and VAP (Fig. 3B, lanes 1 - 3). However, when the eluate containing IncV-3xFLAG was treated with λ phosphatase prior to incubation with GST-VAP_MSP_, the two proteins failed to interact (Fig. 3B, lane 4), indicating that phosphorylation of IncV is necessary for the IncV-VAP interaction *in vitro*.

**Figure 3:**
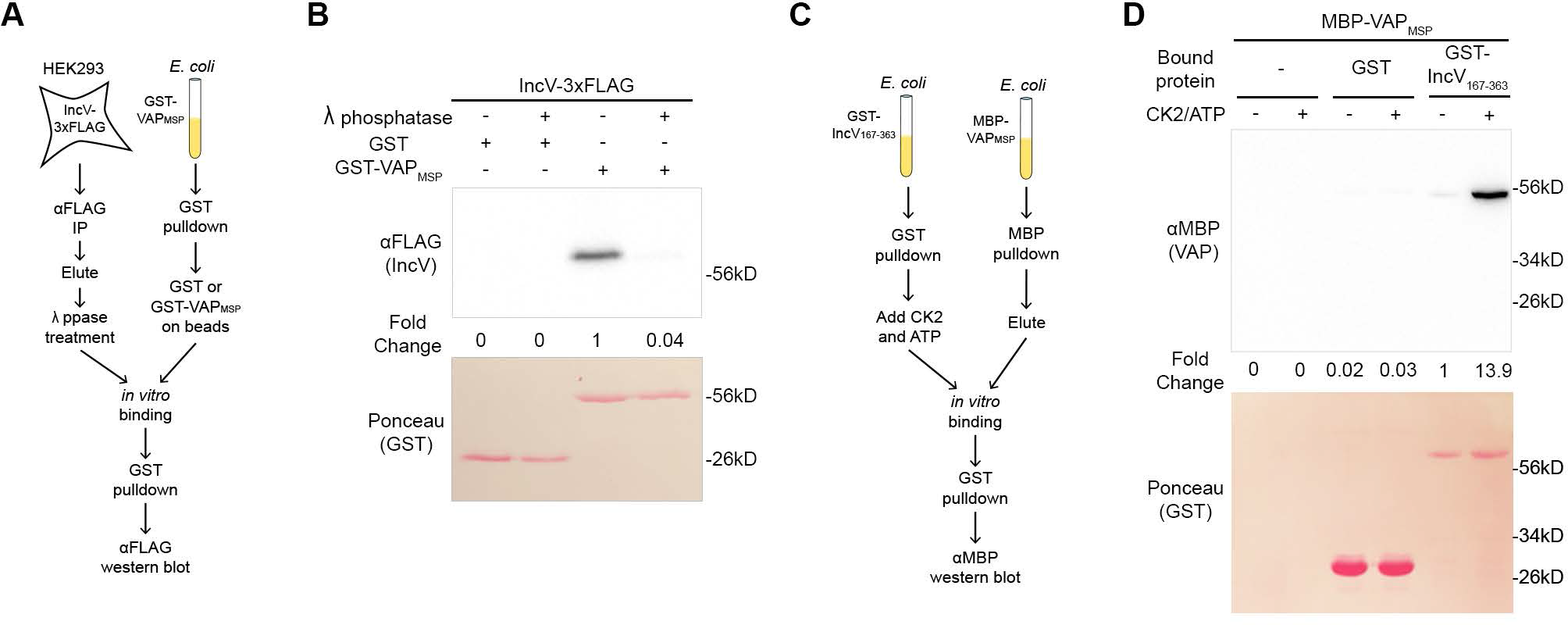
Phosphorylation of IncV is necessary and sufficient to promote the IncV-VAP interaction *in vitro*. (A) Schematic depicting the experimental setup for results in B. (B) *In vitro* binding assay using IncV-3xFLAG purified from HEK293 lysates and treated with lambda (λ) phosphatase (+) or phosphatase buffer alone (-) combined with GST or GST-VAP_MSP_ purified from *E. coli* and immobilized on glutathione beads. The top panel shows proteins detected with anti-FLAG anti-bodies and the bottom panel is the same membrane stained with Ponceau S to detect total protein. (C) Schematic depicting the experimental setup for results in D. (D) *In vitro* binding assay using GST or GST-IncV_167-363_ purified from *E. coli*, and immobilized on glutathione beads, as a substrate for CK2 in the presence (+) or absence (-) of CK2 and ATP, combined with MBP-VAP_MSP_ purified from *E. coli.* The top panel was probed with anti-MBP and the bottom panel was the same membrane stained with Ponceau S to detect the GST construct.

We next determined if IncV phosphorylation by CK2 was sufficient to promote the IncV-VAP interaction in an *in vitro* binding assay (Fig. 3C). MBP-tagged VAP_MSP_ (MBP-VAP_MSP_) and GST- IncV_167-363_ were expressed separately in *E. coli* and purified using amylose resin and glutathione Sepharose beads, respectively. GST-IncV_167-363_ was left attached to glutathione Sepharose beads and was phosphorylated by incubation with recombinant CK2 and ATP before being combined with purified MBP-VAP_MSP_. GST-IncV_167-363_ was pulled down and the samples were subjected to western blot using anti-MBP antibodies (Fig. 3D). Neither the beads alone, nor GST alone pulled down MBP-VAP_MSP_, regardless of whether CK2 and ATP were present or not (Fig. 3D, lanes 1 - 4). In the absence of CK2 and ATP, we observed minimal binding of MBP-VAP_MSP_ to GST- IncV_167-363_ (Fig. 3D, lane 5). However, when GST-IncV_167-363_ was treated with CK2 and ATP prior to GST-pull-down, MBP-VAP_MSP_ and GST-IncV_167-363_ co-immuno-precipitated, indicating that phosphorylation of IncV by CK2 is sufficient to promote the IncV-VAP interaction *in vitro* (Fig. 3D, lane 6). Altogether, these results demonstrate that IncV phosphorylation is necessary and sufficient for the IncV-VAP interaction *in vitro*.

### CK2 is required for IncV phosphorylation and the IncV-VAP interaction during infection

We next determined the contribution of CK2 to IncV phosphorylation and the subsequent assembly of the IncV-VAP tether during *Chlamydia* infection. We first used a genetic approach to deplete CK2β. HeLa cells treated with individual siRNA duplexes targeting *CSNK2B* (A, B, C, or D), or a pool of all four siRNA duplexes (pool), were infected with the *incV* mutant strain of *C. trachomatis* expressing IncV_WT_-3xFLAG from an aTc inducible promoter. The cells were lysed and subjected to western blot analysis. The efficacy of *CSNK2B* knockdown was confirmed by western blot, demonstrating that, in siRNA treated cells, CK2β protein levels ranged from 9.3% to 53.3% compared to control cells (Fig. 4A, middle blot). As shown in Fig. 2A, in control cells, IncV_WT_-3xFLAG appeared as a doublet (Fig. 4A, top blot, left lane, ooo and o). In contrast, depletion of CK2β led to the appearance of additional bands of intermediate apparent molecular weight (Fig. 4A, top and middle blots, pool, A, B, C, D lanes, oo). A line scan analysis of the control sample revealed two peaks corresponding to the top band, corresponding to hyper- phosphorylated IncV (Fig. 4B, black line, left peak, ooo) and to the bottom band, corresponding to unphosphorylated IncV (Fig. 4B, black line, right peak, o). A similar analysis of the banding pattern of IncV upon CK2β depletion, with the pooled or individual siRNA duplexes, revealed the appearance of intermediate peaks between the top and bottom bands, suggesting the formation of hypo-phosphorylated species of IncV (Fig. 4B, middle peaks, oo). These results provided a first indication that CK2 mediates IncV phosphorylation during infection. However, none of the siRNA duplex treatments led to a complete dephosphorylation of IncV, which could be due to the incomplete knockdown of CK2β (Fig. 4A, middle blot).

**Figure 4:**
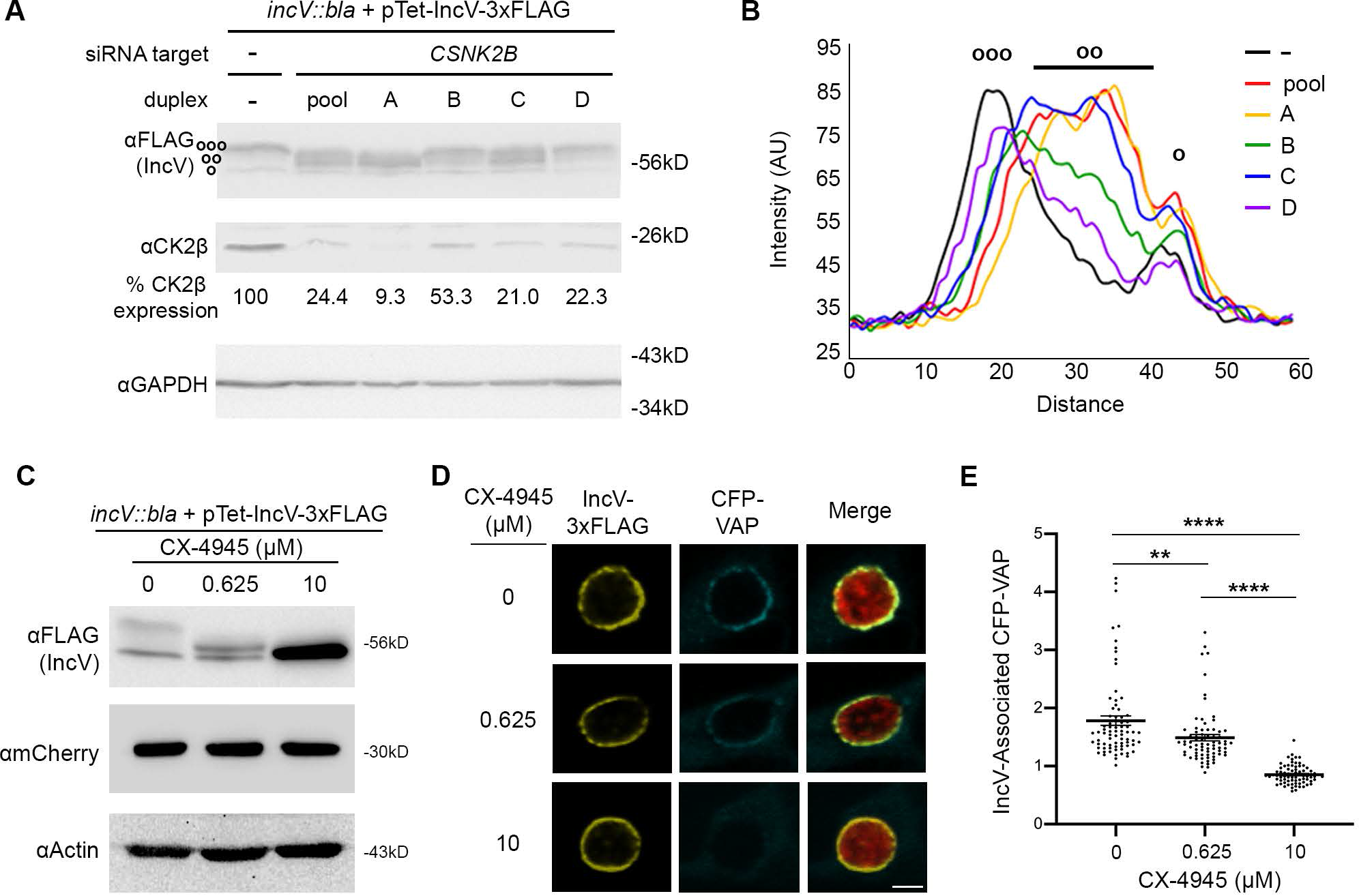
CK2 plays a role in the IncV-VAP interaction during infection. (A) Western blot of lysates of HeLa cells treated with siRNA buffer alone (-) or with siRNA duplexes targeting *CSNK2B* (pool of 4 duplexes or individual duplexes A, B, C, or D) and infected with a *C.trachomatis incV* mutant expressing IncV-3xFLAG. The top panel was probed with anti-FLAG. The middle panel was probed with anti-CK2β. The bottom panel was probed with anti-GAPDH. Relative expression levels of CK2β normalized to GAPDH loading controls are shown as a percentage of no siRNA control expression. (ooo) hyperphosphorylated IncV, (oo) intermediate hypophosphorylated IncV, (o) unphosphorylated IncV. (B) Line Scan analysis of FLAG signal detected in A. The peak on the left (ooo) corresponds to the hyperphosphorylated species of IncV, and the peak on the right (o) corresponds to the unphosphorylated species of IncV. Intermediate hypophosphorylated species are indicated by any peak between the left and right peaks (oo). Each line represents a different condition: Control, black; siRNA pool of duplexes A-D, red; siRNA duplex A, yellow; siRNA duplex B, green; siRNA duplex C, blue; siRNA duplex D, purple. (C-E) HeLa cells, expressing CFP-VAP (D-E only), were infected with *C. trachomatis incV* mutant expressing IncV-3xFLAG under the control of the aTc inducible promoter and treated with increasing concentrations of the CK2 inhibitor CX-4945 (0, 0.625, 10 μM) for two hours at 18 h post infection and prior to the induction of IncV-3xFLAG expression at 20 h post infection. The samples were processed 24 h post infection for western blot (C) or confocal microscopy (D-E). (C) Cell lysates were probed with anti-FLAG (top blot), anti-mCherry (middle blot), or anti-actin (bottom blot) antibodies. (D) Single plane confocal micrographs of HeLa cells expressing CFP- VAPA (blue), infected with *incV* mutant expressing IncV-3xFLAG (yellow) and mCherry (red). The merge is shown on the right. Scale bar is 5μm. (E) Quantification of the mean intensity of the CFP-VAP signal within an object generated from the IncV-3xFLAG signal and normalized to the mean intensity of CFP-VAP in the ER. Each dot represents one inclusion. Data show the mean and SEM of a combination of three independent experiments. One-way ANOVA and Tukey’s post hoc test was performed. ** *P* <0.01, **** *P* <0.0001.

To complement the genetic approach described above, we conducted a pharmacological approach using the CK2-specific inhibitor CX-4945 (Rusin et al., 2017). HeLa cells infected with a *C. trachomatis incV* mutant expressing IncV_WT_-3xFLAG under the control of the aTc inducible promoter were treated with increasing concentrations of CX-4945 (0, 0.625, 10 μM) at 18 h post infection, prior to the induction of IncV_WT_-3xFLAG expression at 20 h post infection. This experimental set up allowed for CK2 inhibition, prior to IncV_WT_-3xFLAG synthesis, translocation, insertion into the inclusion membrane and exposure to the host cell cytosol. The cells were lysed 24 h post infection and subjected to western blot analysis to determine the effect of CK2 inhibition on the apparent molecular weight of IncV. The apparent molecular weight of IncV decreased in a dose-dependent manner (Fig. 4C, top blot), leading to an apparent molecular weight corresponding to unphosphorylated IncV at the 10 μM concentration. These results demonstrate that CK2 activity is essential for IncV phosphorylation during infection.

We next determined whether inhibition of CK2 affected the IncV-dependent VAP recruitment to the inclusion and, therefore, the assembly of the IncV-VAP tether. We used the same experimental setup as above, except that cells expressed CFP-VAP. At 24 h post infection, the cells were fixed, immuno-stained with anti-FLAG antibody, and processed for confocal microscopy. Qualitative and quantitative assessment of the micrographs indicated that CX-4945 did not interfere with IncV translocation and insertion into the inclusion membrane (Fig. 4D and Fig. S1B). As previously observed (Stanhope et al., 2017), IncV_WT_-3xFLAG expression correlated with a strong CFP-VAP association with the inclusion (Fig. 4D, top panels). In comparison, pre-treatment of the cells with 10µM of CX-4945 abolished VAP recruitment to the inclusion (Fig. 4D, bottom panels). Quantification of the CFP-VAP signal associated with IncV at the inclusion membrane confirmed the qualitative analysis and also revealed an intermediate phenotype for cells treated with 0.625µM of CX-4945 (Fig. 4D-E). Altogether, these results demonstrate that phosphorylation of IncV by CK2 is required for the IncV-dependent VAP recruitment to the inclusion.

### Three serine residues in a C-terminal domain of IncV mediate CK2 and VAP recruitment to the inclusion and IncV phosphorylation

To gain further mechanistic insight about the CK2-IncV-VAP interplay, we next determined which domain of IncV was important for the recruitment of CK2 to the inclusion by generating a series of C-terminal truncated IncV constructs (Fig. 5A). These constructs, as well as the full length IncV (FL, 1-363), were cloned under the aTc inducible promoter and expressed from a *C. trachomatis incV* mutant strain. All IncV constructs similarly localized to the inclusion membrane (Fig. S1C). HeLa cells expressing YFP-CK2β were infected with each of the complemented strains, and the ability of the truncated versions of IncV to recruit YFP-CK2β to the inclusion was assessed by confocal microscopy. Qualitative and quantitative analysis revealed that, compared to full length IncV_FL_-3xFLAG, IncV_1-341_-3xFLAG was no longer capable of recruiting YFP-CK2β to the inclusion, whereas IncV_1-356_-3xFLAG was moderately affected (Fig. S3A-B). Additionally, strains expressing IncV_1-341_-3xFLAG also exhibited a significant reduction in IncV-associated VAP compared to IncV_FL_- or IncV_1-356_-3xFLAG (Fig. S3C-D). Altogether, these results demonstrate that a C-terminal region of IncV, between amino acids 342 and 356, is required for the IncV- dependent CK2 recruitment to the inclusion and subsequent VAP association with the inclusion.

**Figure 5:**
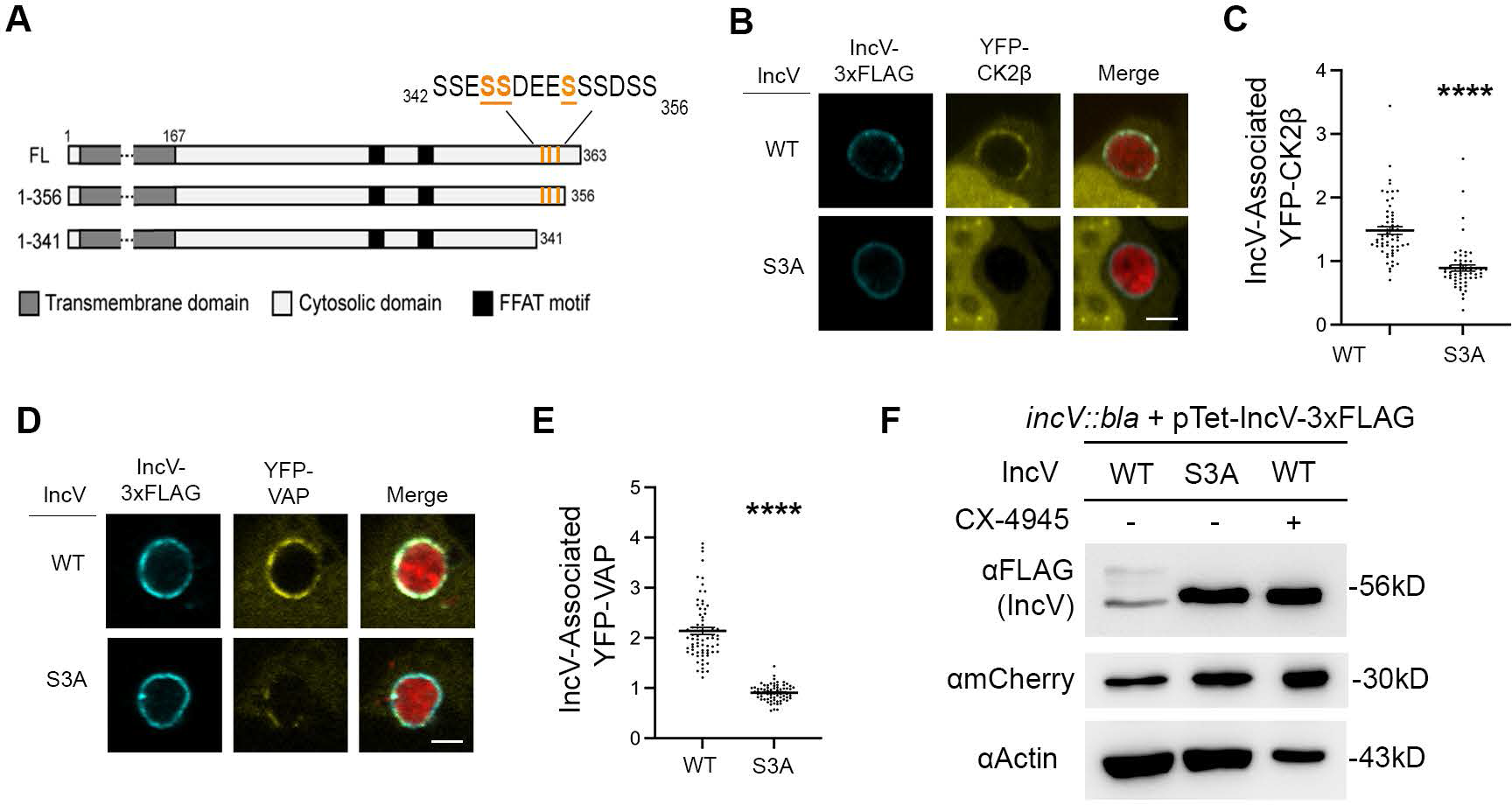
Three serine residues in a C-terminal domain of IncV mediate CK2 and VAP recruitment to the inclusion, and IncV phosphorylation. (A) Schematic depicting truncated IncV constructs. The numbers indicate the amino acid position within the IncV protein sequence. CK2 phosphorylation sites that do not require priming are indicated in orange. (B, D) Single plane confocal images of HeLa cells expressing YFP-CK2β (B) or YFP-VAP (D) (yellow), infected with a *C. trachomatis incV* mutant expressing mCherry constitutively (red) and IncV_WT_-3xFLAG (WT) or IncV_S345A/S346A/S350A_-3xFLAG (S3A) (blue) under the control of the aTc inducible promoter. The merge is shown on the right. Scale bar is 5μm. (C, E) Quantification of the mean intensity of YFP- CK2β (C) and YFP-VAP (E) within the IncV object normalized to the mean intensity of YFP- CK2β in the cytosol and YFP-VAP in the ER, respectively. Data show the mean and SEM of a combination of three independent experiments. \*\*\*\**P* < 0.0001 (Student’s t-test). (F) Western blot of lysates of HeLa cells infected with a *C. trachomatis incV* mutant expressing IncV_WT_-3xFLAG (WT), IncV_S3A_-3xFLAG (S3A), or *C. trachomatis* expressing IncV_WT_-3xFLAG treated with 10 μM CX-4945 as described in Fig. 3C (WT; CX-4945 +) and probed with anti-FLAG (top blot), anti-mCherry (middle blot), and anti-actin (bottom blot) antibodies.

Interestingly, the primary amino acid structure of the IncV domain necessary for CK2 recruitment (_342_SSESSDEESSSDSS_356_) contains seven CK2 recognition motifs (S/T-x-x-D/E/pS/pY) (Litchfield, 2003) (Fig. 5A). Three of them do not require priming by phosphorylation of the fourth serine or tyrosine residue and could result in the direct CK2-dependent phosphorylation of IncV on serine residues S_345_, S_346_, and S_350_, hereby facilitating the assembly of the IncV-VAP tether. To test this hypothesis, all three serine residues were substituted for unphosphorylatable alanine residues (IncV_S345A-S346A-S350A_ referred to as IncV_S3A_). HeLa cells expressing YFP-CK2β or YFP- VAP were infected with *C. trachomatis incV* mutant strains expressing IncV_WT_- or IncV_S3A_- 3xFLAG. The cells were fixed at 24 h post infection, immunostained with anti-FLAG antibody, and analyzed by confocal immunofluorescence microscopy. IncV_WT_- and IncV_S3A_-3xFLAG displayed similar inclusion localization (Fig S1D). However, qualitative and quantitative analysis revealed that in comparison to IncV_WT_-3xFLAG, IncV_S3A_-3xFLAG expression resulted in a significant decrease in both CK2β and VAP recruitment to the inclusion (Fig. 5B-E). Altogether, these results indicate that serine residues S_345_, S_346_, and S_350_ located in a C-terminal motif of IncV, are critical for CK2 recruitment to the inclusion and the CK2-dependent assembly of the IncV- VAP tether.

To determine if IncV_S3A_ failed to interact with VAP because of a lack of IncV phosphorylation, we assessed IncV_S3A_ apparent molecular weight by western blot analysis of lysates from HeLa cells infected with a *C. trachomatis incV* mutant expressing IncV_WT_- or IncV_S3A_-3xFLAG. Compared to IncV_WT_-3xFLAG, which as previously observed ran as a doublet corresponding to both phosphorylated and unphosphorylated species of IncV (Fig. 5F, lane 1), the apparent molecular weight of IncV_S3A_-3xFLAG (Fig. 5F, lane 2), was identical to that of unphosphorylated IncV_WT_-3xFLAG upon treatment with the CK2 inhibitor CX-4945 (Fig. 5F, lane 3). These results indicated that IncV_S3A_ is unphosphorylated and suggested that phosphorylation of S_345_, S_346_, and S_350_ may be sufficient to mediate the *in vitro* IncV-VAP interaction observed upon CK2 phosphorylation of IncV (Fig. 3D). To test this, S_345_, S_346_, and S_350_ were substituted to for phosphomimetic aspartic acid residues. The corresponding IncV construct, referred to as IncV_S3D_, was purified from *E. coli* and tested for VAP binding *in vitro*. IncV_S3D_ did not result in a significant increase in VAP binding compared to IncV_WT_ (Fig. S4). Altogether, these results indicate that, although critical for CK2 recruitment, assembly of the IncV-VAP tether at the inclusion, and IncV phosphorylation status, phosphorylation of S_345_, S_346_, and S_350_ alone is not sufficient to promote VAP binding *in vitro*, suggesting that additional IncV phosphorylation sites are required to promote optimal interaction between IncV and VAP.

### Phosphorylation of serine rich tracts upstream of IncV FFAT motifs substitute typical acidic tracts and are key for the IncV-VAP interaction

In addition to the seven amino acid core of the FFAT motif, VAP-FFAT mediated interactions also rely on the presence of acidic residues upstream of the core sequence, referred to as the acidic tract. It allows for the initial electrostatic interaction with VAP by interacting with the electropositive charge of the MSP domain before the FFAT core motif locks into its dedicated groove (Furuita et al., 2010). We noted that, instead of typical acidic residues, the primary amino acid structures upstream of the IncV FFAT motifs are highly enriched in phosphorylatable serine residues (Fig. 6A). We hypothesized that, if phosphorylated, these serine residues could serve as an acidic tract and facilitate the IncV-VAP interaction. To test this hypothesis, the 10 residues directly upstream of the phospho-FFAT motif and the 8 residues directly upstream of the canonical FFAT motif were mutated to alanine residues (referred to as IncV_S/A_) and the ability of IncV_S/A_- 3xFLAG to recruit VAP to the inclusion was assessed. HeLa cells expressing YFP-VAP were infected with *C. trachomatis incV* mutant strains expressing either IncV_WT_-, IncV_F263A/Y287A_-, or IncV_S/A_-3xFLAG under an aTc inducible promoter. The cells were fixed at 24 h post infection and analyzed by confocal immunofluorescence microscopy (Fig. 6B). All IncV constructs were equally localized to the inclusion membrane (Fig. S1E). Qualitative and quantitative analysis revealed that expression of IncV_S/A_-3xFLAG resulted in a significant decrease in YFP-VAPA recruitment to the inclusion as observed with IncV_F263A/Y287A_-3xFLAG and compared to IncV_WT_-3xFLAG (Fig. 6B- C). To determine if this decrease in VAP recruitment was due to a lack of CK2 recruitment, the ability of these strains to recruit YFP-CK2β to the inclusion was assessed by confocal microscopy (Fig. S5). All three strains recruited CK2 to the inclusion (Fig. S5), indicating that the lack of VAP recruitment upon expression of IncV_S/A_ was not due to a lack of CK2 recruitment, as observed for IncV_S3A_ (Fig. 5B-E).

**Figure 6:**
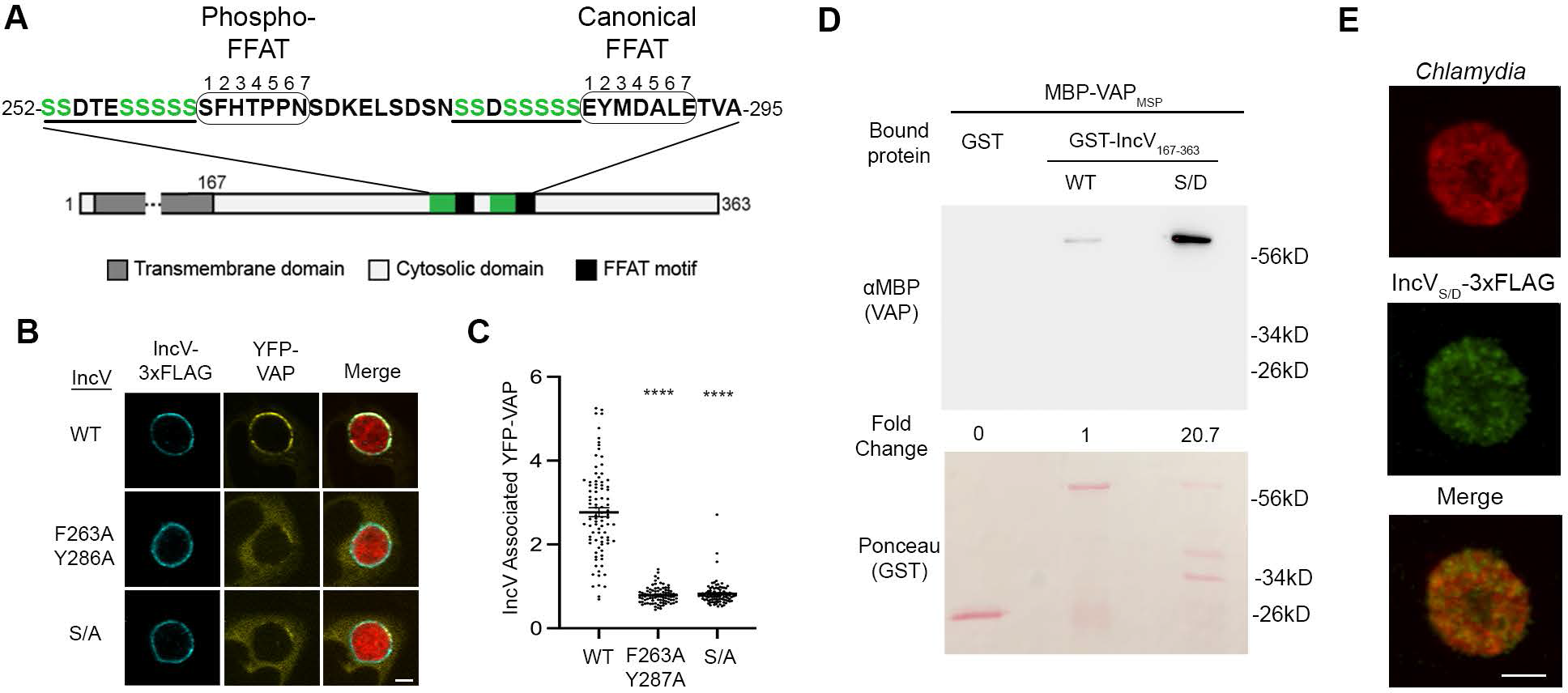
Phosphorylation of the serine tracts upstream of the IncV FFAT motifs facilitates the IncV-VAP interaction. (A) Schematic depicting the IncV protein. The transmembrane domain, the cytosolic domain, and the phospho and canonical FFAT motif cores are indicated in dark grey, light grey, and black, respectively. The amino acid sequence of the FFAT motif cores (Circled) and their respective upstream sequence is shown. The serine-rich tracts are underlined. Serine residues are in green. Numbers 1-7 indicate the amino acid position within the FFAT motif cores, other numbers indicate the amino acid position within the IncV protein sequence. (B) Single plane confocal images of HeLa cells expressing YFP-VAPA (yellow), infected with a *C. trachomatis incV* mutant expressing mCherry consitutively (red) and IncV_WT_-3xFLAG (WT), IncV_F263A/Y287A_-3xFLAG (F263A/Y287A), or IncV_S/A_-3xFLAG (S/A) (blue) under the control of an aTc inducible promoter. The merge is shown on the right. Scale bar is 5μm. (C) Quantification of the mean intensity of YFP-VAP within an object generated from the IncV-3xFLAG signal and normalized to the mean intensity of YFP-VAP in the ER. Each dot represents one inclusion. Data show the mean and SEM of a combination of three independent experiments. One-way ANOVA with Tukey’s post hoc test was performed. **** *P* <0.0001. (D) *In vitro* binding assay using GST, GST-IncV_WT_, or GST-IncV_S/D_ purified from *E. coli*, and immobilized on glutathione beads and combined with MBP-VAP purified from *E. coli.* The top panel was probed with anti-MBP and the bottom panel was the same membrane stained with Ponceau S to detect the GST construct. Note that the IncV and VAP constructs, only include the cytosolic domain of IncV (aa 167-363) and the MSP domain of VAP, respectively. (E) Single plane confocal images of HeLa cells infected with a *C. trachomatis incV* mutant expressing mCherry consitutively (red) and IncV_S/D_-3xFLAG (green) under the control of an aTc inducible promoter in the presence of aTc. The merge is shown on the bottom. Scale bar is 5μm.

We next determined if phosphomimetic mutation of the serine-rich tracts of IncV to aspartic acid residues (referred to as IncV_S/D_) was sufficient to rescue the ability of the cytosolic domain of IncV expressed in *E. coli* to interact with the MSP domain of VAP in our VAP binding *in vitro* assay. As observed before, there was minimal binding of VAP_MSP_ to IncV_WT_ (Fig. 6D, lane 2). However, we observed a 20-fold increase in VAP_MSP_ binding to IncV_S/D_ compared to IncV_WT_ (Fig. 6D, lane 3), indicating that phosphomimetic mutation of the serine-rich tracts is sufficient to promote the IncV-VAP interaction *in vitro*. Altogether, these results indicate that instead of typical acidic tracts, phosphorylated serine-rich tracts located upstream of IncV FFAT motifs are both necessary and sufficient for promoting the IncV-VAP interaction.

In order to confirm the role of IncV serine-rich tracts in promoting the IncV-VAP interaction during infection, we assessed the ability of IncV_S/D_-3xFLAG to recruit VAP to the inclusion, when expressed from an *incV* mutant strain of *C. trachomatis*. In comparison to IncV_WT_ and all other mutated alleles used in this study (Fig. S1), IncV_S/D_-3xFLAG remained trapped within the bacteria and did not localize to the inclusion membrane (Fig. 6E). These results suggest that phosphorylatable serine residues may have been selected over acidic residues to allow proper Type III translocation of IncV to the inclusion membrane.

## Discussion

Based on our results we propose the following model of assembly of the IncV-VAP tether at ER-Inclusion MCS. Unphosphorylated IncV is translocated across the inclusion membrane by the T3SS. Upon insertion into the inclusion membrane and exposure to the cytosol, IncV is phosphorylated by host cell kinases, leading to VAP recruitment and assembly of the IncV-VAP tether. IncV phosphorylation most likely occurs in stages. A first event is the IncV-dependent recruitment of the host kinase CK2 through the C-terminal domain of IncV containing three serine residues that are part of CK2 recognition sites (Fig. 7, Step 1). As a consequence, IncV becomes hyper-phosphorylated, including phosphorylation of T265 of the phospho-FFAT and serine tracts directly upstream of the FFAT motifs (Fig. 7, Step 2). We note that kinases other than CK2 must be involved in this second step, since the phospho-FFAT is not a CK2 target. Phosphorylation of the serine tract and of the phospho-FFAT result in full mimicry of eukaryotic FFAT motifs, leading to IncV interaction with VAP and tether assembly (Fig. 7, Step3). Importantly, the post-translocation phosphorylation of IncV ensure optimal VAP binding while preserving proper T3SS- mediated translocation of IncV to the inclusion membrane. Below we discuss our results in the context of emerging regulatory mechanisms of cellular MCS assembly and highlight conserved and pathogen-specific mechanisms.

**Figure 7:**
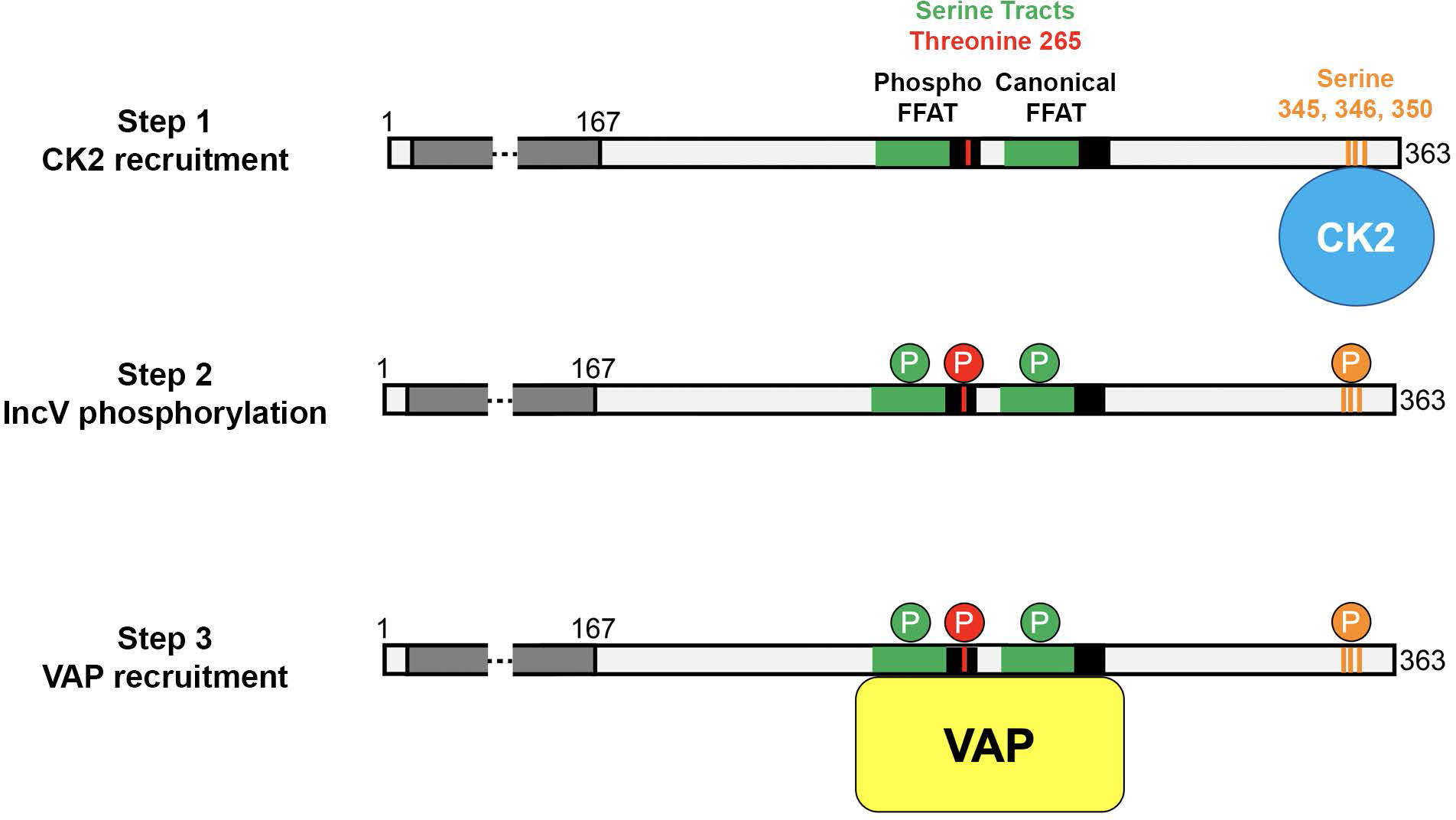
Model of assembly of the IncV-VAP tether of ER-Inclusion MCS. Step 1: After secretion and insertion of unphosphorylated IncV into the inclusion membrane, IncV recruits CK2 (blue) via 3 serine residues S345, S346 and S350 (orange) that are part of CK2 recognition motifs. Step 2: IncV becomes hyperphosphorylated, including phosphorylation of the phospho-FFAT on threonine residue T265 (red) and the serine-rich tract (green) immediately upstream of FFAT core motifs (black). Step 3: IncV phosphorylation leads to full mimicry of FFAT motifs and binding to VAP (yellow). The dark and light grey bars represent the transmembrane and cytosolic domain of IncV, respectively. P represent the phosphorylation of specific residues.

### IncV-dependent recruitment of CK2 to the inclusion

Few kinases phosphorylating VAP-dependent tethers have been identified so far (Xu et al., 2020) and how they associate with MCS to phosphorylate their target has not been explored. Here, we show that IncV recruits CK2 to ER-Inclusion MCS through interaction with its C-terminal domain, a mandatory step for IncV hyper-phosphorylation. Three serine residues (S_345_, S_346_, and S_350_) that match the CK2 recognition motifs (S-x-x-D/E) located in a C-terminal domain of IncV are essential for CK2 recruitment to the inclusion, and IncV phosphorylation. However, while CK2 was required for ER-Inclusion MCS formation during infection, and for IncV-VAP interaction *in vitro*, the introduction of phosphomimetic mutations at S_345_, S_346_, and S_350_ was not sufficient to promote the IncV-VAP interaction *in vitro*, suggesting that additional phosphorylation sites exist. Kinase-substrate recognition is a complex process that goes beyond the simple recognition of a consensus sequence and can involve docking sites away from the phosphorylation sites (Miller & Turk, 2018). The cytosolic domain of IncV contains a large number of additional potential CK2 recognition sites. We therefore propose that alanine substitution of S_345_, S_346_, and S_350_ eliminates an essential docking site for subsequent CK2-mediated phosphorylation of distal residues in the cytosolic domain of IncV, including the serine tracts next to the FFAT motifs (see below). Further investigation of the IncV-dependent recruitment of CK2 to ER-Inclusion MCS could offer some insights into kinase targeting to cellular MCS. Moreover, since intracellular pathogens often mimic cellular processes, our study may have identified CK2 as a regulator of cellular MCS.

### IncV harbors a phospho-FFAT motif

Our results indicate that threonine residue 265 (T_265_) at position 4 of the non-canonical FFAT of IncV is essential for tether assembly. These results experimentally validate the presence of a phospho-FFAT in IncV, as previously proposed (Di Mattia et al., 2020), and add to the growing list of proteins that interact with VAP via a phospho-FFAT. These include STARD3 at ER-endosome contacts, the potassium channel Kv.2 at ER-PM contacts in neurons, and Miga at ERMCS (Di Mattia et al., 2020; Johnson et al., 2018; Xu et al., 2020). Moreover, although not recognized as such at the time, a phospho-FFAT in the norovirus protein NS2 is essential for interaction with VAP and viral replication (McCune et al., 2017), indicating that this mechanism of interaction with VAP is also conserved amongst pathogens.

In the context of the STARD3-dependent formation of ER-endosome contacts, the presence of a single phospho-FFAT is proposed to act as a molecular switch to regulate contact formation (Di Mattia et al., 2020). In addition to a phospho-FFAT, IncV also contains a canonical FFAT, for which, based on current knowledge, the binding to VAP is not subjected to regulation. PTPIP51, an ER-mitochondria contact protein, also contains a combination of a FFAT and a phospho-FFAT (Di Mattia et al., 2018). It is unclear how a most likely constitutive and a regulated FFAT motif cooperate, if one is dominant over the other, and how advantageous such a combination is with respect with MCS regulation. In the case of IncV, one could speculate that the canonical FFAT motif allows for a baseline level of VAP recruitment to the inclusion and MCS formation while the phospho-FFAT allows for the increase in VAP recruitment beyond this baseline. We note that T265 is not a CK2 target, and therefore in addition to CK2, at least one additional kinase must be involved in IncV phosphorylation.

### IncV-VAP interaction is mediated by phosphorylatable serine tracts

In eukaryotic FFAT motifs, a number of negative charges upstream of the FFAT motif is proposed to facilitate the initial interaction with the MSP domain of VAP (Furuita et al., 2010). This electronegative surface is conferred by acidic residues, but phosphorylated residues have been implicated in two instances. The phosphorylation, by an unknown kinase, of a single serine residue six amino acids upstream of the CERT FFAT motif (S315) enhances the CERT-VAP interaction (Kumagai et al., 2014), while six serine residues, spread over 21 residues upstream of the core FFAT motif of Miga, facilitate the Miga-VAP interaction (Xu et al., 2020). At least two kinases CKI and CaMKII, were required for Miga phosphorylation; however, other kinases are likely involved (Xu et al., 2020). In the case of IncV, the mimicry of an acidic track via phosphorylatable residues seems to be brought to the extreme, since the eight to ten amino acid stretch directly preceding each FFAT motif include 80 to 87% of serine residues, the remaining residues being acidic. Except for OSBP2/ORP4, which contains 6 acidic residues (including a phosphorylatable threonine), most acidic tracts contain few acidic residues directly upstream of the core of the FFAT motif (Neefjes & Cabukusta, 2021). IncV is the first example of a FFAT motif-containing protein that displays a serine tract in place of acidic tract. If built into the available FFAT motif identification algorithms, this feature could potentially reveal additional cellular VAP interacting proteins (Di Mattia et al., 2020; Murphy & Levine, 2016).

### Phospho-regulation and pathogenesis

During co-evolution with the mammalian host, obligate intracellular bacteria such as *Chlamydia* have evolved to take advantage of and manipulate cellular machinery. One mechanism is *via* molecular mimicry, in which the pathogen mimics features that are uniquely present in host proteins (Mondino et al., 2020). In the case of IncV and acidic tracks in FFAT motifs, however, one could wonder why evolution would converge toward a mechanism relying on phosphorylation by host cell kinases, as opposed to simply selecting for genetically encoded acidic residues. In the case of *Chlamydia* Inc proteins, it is possible that tracks of aspartic acid or glutamic acid residues would create an excess of negative charges that may interfere with Type III secretion. In support of this notion, we found that *Chlamydia* IncV is no longer properly translocated to the inclusion membrane when the serine tracts are mutated to aspartic acid residues, and instead remains trapped within the bacteria. Our results support the notion that the recruitment of CK2 to the inclusion supports the assembly of the IncV-VAP tether. In addition, we cannot exclude the possibility that the recruitment of a phosphatase to ER-Inclusion MCS may contribute to the disassembly of IncV- VAP tethers, as shown for the calcineurin-dependent disassembly of Kv.2-VAP ER-PM contacts in neurons (Park et al., 2006). A combination of host cell kinases and phosphatases could thus regulate the dynamics of ER-Inclusion contact sites during the *Chlamydia* developmental cycle.

## Materials and Methods

### Ethics statement

All genetic manipulations and containment work were approved by the UVA Biosafety Committee and are in compliance with the section III-D-1-a of the National Institutes of Health guidelines for research involving recombinant DNA molecules.

### Cell lines and bacterial strains

HeLa cells (ATCC CCL-2) and HEK293 cells (ATCC CRL-1573) were maintained in DMEM high glucose (Gibco) containing 10% heat-inactivated fetal bovine serum (Gibco) at 37°C and 5% CO_2_. *Chlamydia trachomatis* Lymphogranuloma venereum, type II (ATCC L2/434/Bu VR-902B) was propagated in HeLa cells as previously described (Derré et al., 2007). The *incV::bla* mutant strain of *C. trachomatis* (also known as *CT005::bla*) was obtained from Ted Hackstadt (NIH, Rocky Mountain Laboratories) (Weber et al., 2017).

### Plasmid construction

Plasmids were constructed using the primers (IDT) and templates listed in Table S1, Herculase DNA polymerase (Stratagene), restriction enzymes (NEB), and T4 DNA ligase (NEB).

### Vectors for expression in mammalian cells

The IncV-3xFLAG construct cloned in the pCMV-IE-N2-3xFLAG vector was previously described (Stanhope et al., 2017). The YFP-CK2α and YFP-CK2β plasmids were kind gifts from Claude Cochet and Odile Filhol-Cochet (Institut Albert Bonniot Departement Reponse et Dynamique Cellulaires) and were previously characterized (Filhol et al., 2003). The CFP-VAP and YFP-VAP plasmids were constructed by cloning the VAPA open reading frame (ORF) into pCMV-N1-CFP and pCMV-N1-YFP, respectively, using AgeI and HindIII restriction sites.

### Vectors for expression in *E. coli*

The GST-VAP_MSP_ plasmid was previously described (Stanhope et al., 2017). MBP-VAP_MSP_ was constructed by cloning the MSP domain of VAPA using NotI and BamHI into pMAL. The GST- IncV_167-363_ fusion constructs WT, S3D, or S/D were generated by cloning a DNA fragment encoding amino acids 167-363 of IncV into the BamHI and XhoI restriction sites of pGEX-KG. \

### Vectors for expression in *C. trachomatis*

Full length, truncated versions, and mutant versions of IncV were cloned into the p2TK2_Spec_-SW2 mCh(Gro) vector as previously described (Cortina et al., 2019). Briefly, TetRTetAP promoter and 3xFLAG *incD* terminator fragments were appended onto either end of the full length IncV fragment using overlap PCR to generate TetRTetAP-IncV-3xFLAG-IncDterm fragments (Tet- IncV-3xFLAG for short). Truncated IncV constructs (DNA corresponding to amino acids 1-341 or 1-356) were generated using overlap PCR to truncate the IncV ORF. Mutant IncV constructs were generated using overlap PCR to substitute amino acids of the IncV ORF. All versions of Tet- IncV-3xFLAG were cloned into p2TK2Spec-SW2 mCh(gro) using KpnI and NotI. mCherry is expressed constitutively from the *groESL* promoter, and IncV-3xFLAG variants (T265A, T265A/Y287A, Full length, 1-356, 1-341, S3A, S/A, and S/D) are expressed under the aTc-inducible promoter. The IncV_F263A/Y287A,_ and IncV_Y287A_ plasmids were previously described (Stanhope et al., 2017).

### *C. trachomatis* transformation and *incV::bla* complementation

Wild type *C. trachomatis* or an *incV* mutant (*incV::bla*) were transformed with pTet-IncV_WT_- 3xFLAG, pTet-IncV_Y287A_-3xFLAG, pTet-IncV_F236A/Y287A_-3xFLAG, pTet-IncV_T265A_-3xFLAG, pTet-IncV_T265A/Y287A_-3xFLAG, pTet-IncV_1-356_-3xFLAG, pTet-IncV_1-341_-3xFLAG, pTet-IncV_S3A_-3xFLAG, pTet-IncV_S/A_-3xFLAG, or pTet-IncV_S/D_-3xFLAG using our previously described calcium-based *Chlamydia* transformation procedure (Cortina et al., 2019).

### DNA transfection

Cells were transfected with mammalian construct DNA according to manufacturer instructions with X-tremeGENE 9 DNA Transfection Reagent (Roche).

### SDS-PAGE

Cells were either directly lysed in 2x Laemmli buffer with 10mM DTT or IncV was purified as described in the immunoprecipitation and protein purification sections then suspended in a final concentration of 1x Laemmli buffer with 10mM DTT. Protein samples were separated using SDS- PAGE.

### Immunoblotting

After SDS/PAGE, proteins were transferred onto nitrocellulose membranes (GE Healthsciences). Prior to blocking, membranes were stained with Ponceau S in 5% acetic acid and washed in dH_2_O. Membranes were incubated for 1 hour with shaking at room temperature in blocking buffer (5% skim milk in 1x PBS with 0.05% Tween). Membranes were then incubated with primary and secondary (HRP-conjugated) antibodies diluted in blocking buffer overnight at 4°C and 1 hour at room temperature, respectively, with shaking. ECL Standard western blotting detection reagents (Amersham) were used to detect HRP-conjugated secondary antibodies on a BioRad ChemiDoc imaging system. CK2β was detected using secondary antibodies conjugated to Alexa Fluor 800 on Li-Cor Odyssey imaging system.

### Antibodies

The following antibodies were used for immunofluorescence microscopy (IF) and immunoblotting (WB): mouse monoclonal anti-FLAG [1:1,000 (IF); 1:10,000 (WB); Sigma], rabbit polyclonal anti-CK2β [1:200 (IF); 1:1,000 (WB); Bethyl Antibodies]; rabbit polyclonal anti-thiophosphate ester antibody [1:2000 (WB); Abcam], rabbit polyclonal anti-MBP [1:10,000 (WB); NEB], rabbit polyclonal anti-GAPDH [1:10,000 (WB); ], rabbit polyclonal anti-mCherry [1:2,000 (WB); BioVision], rabbit polyclonal anti-actin [1:10,000 (WB); Sigma], HRP-conjugated goat anti-rabbit IgG [1:10,000 (WB); Jackson], HRP-conjugated goat anti-mouse IgG [1:10,000 (WB); Jackson], Alexa Fluor 514-, 800-, or Pacific Blue-conjugated goat anti-mouse IgG [1:500 (IF); 1:10,000 (WB); Molecular Probes].

### Immunoprecipitation of IncV-3xFLAG from HEK293 cells infected with C*. trachomatis*

800,000 HEK293 cells were seeded into one well of a six-well plate (Falcon) and infected the following day with *C. trachomatis* at a multiplicity of infection (MOI) of 5. 8 hours post infection, media containing 2ng/mL anhydrotetracycline (aTc) was added to the infected cells for 16 hours. 24 hours post-infection, culture media was removed from the cells and 500μL of lysis buffer (20 mM Tris pH 7.5, 150 mM NaCl, 2 mM EDTA, 1% Triton X-100, protease inhibitor mixture EDTA-free (Roche)) was added per well. Cells were lysed for 20 minutes at 4°C with rotation. Lysates were centrifuged at 16,000xg for 10 minutes at 4°C to pellet nuclei and unlysed cells. Cleared lysates were incubated with 10μL of anti-FLAG M2 affinity beads (Sigma) for 2 hours at 4°C with rotation. The beads were washed with lysis buffer three times. Proteins were eluted with 50μL of 100μg/mL 3xFLAG peptide (Sigma) in 1x Tris-buffered saline. For cells transfected with pCMV-IE-N2-IncV-3xFLAG, cells were not infected, and the remainder of the protocol remained the same starting with removal of media and lysing.

### Phosphatase assay

Immunoprecipitation was performed as described above with the following changes: The beads were washed with 1x Tris-buffered saline (TBS) three times and proteins were eluted with 55μL of 100μg/mL 3xFLAG peptide (Sigma) in 1x TBS. 20μL of eluate was combined with 2.5μL of 10mM MnCl2, 2.5μL of 10x PMP buffer (NEB), and 400 units of lambda (λ) phosphatase (NEB) for 24 hours at 4°C. The assay was halted by adding 5μL of 6x Laemmli buffer with 10mM DTT. Samples were boiled and 10μL of sample was then used in SDS-PAGE.

### DNA transfections and infections for microscopy

HeLa cells were seeded onto glass coverslips and transfected with YFP-CK2 (α or β), CFP-VAPA, or YFP-VAPA the following day. 24 hours post-transfection, cells were infected with the indicated strain of *C. trachomatis* at a MOI of 1. 20 hours post-infection, media containing 20ng/mL aTc (final concentration) was added for 4 hours to induce expression of IncV-3xFLAG.

### Immunofluorescence and confocal microscopy

HeLa cells seeded on glass coverslips and infected with *C. trachomatis* were fixed 24 hours post-infection with 4% paraformaldehyde in 1x PBS for 20 minutes at room temperature then washed with 1x PBS three times. The coverslips were sequentially incubated with primary and secondary antibodies in 0.1% Triton X-100 in 1x PBS for 1 hour at room temperature. For coverslips stained with anti-CK2β, antibodies were diluted in 0.5% Triton X-100 and 5% BSA in 1x PBS. Coverslips were washed with 1x PBS three times then mounted with glycerol containing DABCO and Tris pH 8.0. Confocal images were obtained using an Andor iXon ULTRA 888BV EMCCD camera and a Yokogawa CSU-W1 Confocal Scanner Unit attached to a Leica DMi8 microscope. 1 μm thick Z slices covering the entirety of the cell were captured.

### Quantification of YFP-CK2β, CFP-VAP, and YFP-VAP inclusion association

Quantification of YFP-CK2β, CFP-VAP, or YFP-VAP association with IncV-3xFLAG on the inclusion membrane was performed using the Imaris imaging software. First, three-dimensional (3D) objects were generated from the raw signal of IncV-3xFLAG on the inclusion membrane. Objects were edited such that IncV-3xFLAG colocalizing with the mCherry bacteria was removed. Within the resulting IncV object, the mean intensity of YFP-CK2β, CFP-VAP, or YFP-VAP was calculated by the Imaris software and normalized to the mean intensity of YFP-CK2β, CFP-VAP, or YFP-VAP within the cytosol surrounding the inclusion.

Quantification of the IncV-3xFLAG volume was performed to ensure there was no defect in inclusion localization. Using the Imaris imaging software, the sum of the voxels corresponding to the IncV-3xFLAG signal above the threshold set by the signal within the cytosol was calculated for IncV-3xFLAG and mCherry. The IncV-3xFLAG volume was normalized to its corresponding inclusion volume.

Each experiment was performed in triplicate with at least 20-30 inclusions analyzed per condition per replicate. Unless specified, data from 3 independent replicates are combined into a single graph. Each point on the graph represents a single inclusion with the average value and SEM shown. Student’s t-tests or one-way ANOVA with multiple comparisons were performed.

### Protein purification

Expression of GST, GST-VAP_MSP_, GST-IncV_167-363_, GST-IncV_167-363_ S/D or MBP-VAP_MSP_ was induced for two hours by the addition of isopropyl-β-ᴅ-thiogalactopyranoside (0.1mM, final concentration) to a 10 mL culture of *E. coli* BL21-λDE3 at OD 0.8. Bacterial pellets were stored at -80°C. Frozen pellets were thawed and resuspended in 800μL sonication buffer (20 mM Tris pH 7.5, 300 mM NaCl, 2 mM EDTA, 1 mM MgCl2, 1% Triton X-100, 1mM DTT, 1mM PMSF). The samples were sonicated using five 5-second pulses at 40% power then centrifuged at 13,000xg for 10 minutes at 4°C. 40μL of glutathione Sepharose beads (GE) for GST-tagged constructs and 40μL of Amylose resin for MBP-tagged constructs were washed three times with sonication buffer then added to the cleared lysate and incubated for 2 hours at 4°C with rotation. The beads were washed three times in TBS.

### *In vitro* kinase assay

Protein bound glutathione Sepharose beads were resuspended in 1x NEBuffer™ for Protein Kinases supplemented with 1mM ATPγS and 10 units of CK2 (NEB) and incubated at 30°C for 45 minutes. P-Nitrobenzyl mesylate (PNBM) was added to the kinase reaction at a final concentration of 2.4mM for 2 hours at room temperature in the dark. The PNBM alkylation reaction was quenched by adding an equal volume of 2x Laemmli buffer. Proteins were separated using SDS-PAGE on a 12% acrylamide gel then transferred to a nitrocellulose membrane. The membrane was stained with Ponceau S in 5% acetic acid to detect total protein then washed in dH_2_O. The membrane was then probed with anti-thiophosphate ester antibodies to detect phosphorylated proteins which were detected with HRP-conjugated secondary antibodies.

### *In vitro* binding assay

First, GST, GST-IncV_167-363_, GST-IncV_167-363_ S/D, and MBP-VAP_MSP_ were purified as described in protein purification. MBP-VAP_MSP_ was eluted from amylose resin using 100μL 1x TBS supplemented with 10mM maltose monohydrate. GST, GST-IncV_167-363_, or GST-IncV_167-363_ S/D attached to glutathione beads were washed three times in sonication buffer. 500μL of sonication buffer containing 1.25μg MBP-VAP_MSP_ was added to each tube with GST beads and binding was allowed to occur overnight at 4°C with rotation. Following overnight binding, beads were washed three times in 1x TBS. After the final wash, all liquid was removed from the beads which were then suspended in 20μL 2x Laemmli buffer. The entire sample was separated by SDS-PAGE, proteins transferred to a nitrocellulose membrane which was stained with Ponceau S to detect the GST construct then probed with anti-MBP to detect MBP-VAP_MSP_.

### *In vitro* binding assay with IncV dephosphorylation

First, the phosphatase assay was performed with the following changes: 1,000,000 HEK293 cells stably transfected with pCMV-IE-N2-IncV-3xFLAG were seeded per 6 well. 6 wells were lysed in 500μL lysis buffer each and lysates from two wells were combined. 10μL of anti-FLAG beads were added per 1000μL cleared lysate for 2 hours at 4°C with rotation. All beads were combined after the first wash, and proteins were eluted in 150μL elution buffer (130μL eluate collected).

Next, GST and GST-VAP_MSP_ were purified as described in protein purification. Per phosphatase assay tube: 1.5μg of GST or GST-VAP_MSP_ attached to beads (determined empirically by comparison of Coomassie stained gel to BSA standard curve) were suspended in 500μL lysis buffer then added to tubes containing the IncV-3xFLAG-containing eluate (+/- phosphatase treatment). Binding was allowed to occur overnight at 4°C with rotation.

To confirm that IncV dephosphorylation was successful, a set of control tubes were incubated with beads alone (no GST construct) in lysis buffer to mimic experimental conditions.

24 hours after binding, beads were washed three times in 1x TBS. After the final wash, all liquid was removed from the beads which were then suspended in 20μL 2x Laemmli buffer. The entire sample was separated by SDS-PAGE, proteins transferred to a nitrocellulose membrane which was stained with Ponceau S to detect the GST construct then probed with anti-FLAG to detect IncV- 3xFLAG.

### *In vitro* binding assay with CK2 phosphorylation of IncV

First, GST, GST-IncV_167-363_, and MBP-VAP_MSP_ were purified as described in protein purification. MBP-VAP_MSP_ was eluted from amylose resin using 100μL 1x TBS supplemented with 10mM maltose monohydrate. 1.5μg of GST or GST-IncV_167-363_ attached to glutathione beads (determined empirically by comparison of Coomassie stained gel to BSA standard curve) or beads alone were suspended in 1x NEBuffer™ for Protein Kinases with 200μM ATP (Thermo) and 100 units of CK2 (NEB) at 30°C for 45 minutes. Beads were washed three times in sonication buffer. 1.25μg MBP-VAP_MSP_ suspended in 500μL sonication buffer was added to each tube with beads and binding was allowed to occur overnight at 4°C with rotation. 24 hours after binding, beads were washed three times in 1x TBS. After the final wash, all liquid was removed from the beads which were then suspended in 20μL 2x Laemmli buffer. The entire sample was separated by SDS-PAGE, proteins transferred to a nitrocellulose membrane which was stained with Ponceau S to detect the GST construct then probed with anti-MBP to detect MBP-VAP_MSP_.

### GST-Pull down immunoblot quantification

Immunoblots and Ponceau S staining were quantified using the ImageJ software (NIH). The immunoblot band intensity was normalized to the Ponceau S band intensity and the fold change determined relative to wild type or untreated conditions.

### CK2 depletion using siRNA

CK2 was depleted from cells using a pool of four siRNA duplexes or each duplex individually that was transfected with Dhamafect 1 transfection reagents. On day 0, one volume of 200nM siRNA in siRNA buffer was incubated with one volume of 5μL/mL of Dharmafect 1 transfection reagent in DMEM high glucose in a well for 20 minutes at room temperature. Two volumes of DMEM High Glucose supplemented with 20% FBS and 200,000 HeLa cells per mL were added to the well. Cells were incubated at 37°C with 5% CO_2_ for three days. The total volume for one 96 well was 120μL. The *CSNK2B* target sequence for each individual siRNA duplex was: A, CAACCAGAGUGACCUGAUU; B, GACAAGCUCUAGACAUGAU; C, CAGCCGAGAUGCUUUAUGG; D, GCUCUACGGUUUCAAGAUC. The efficacy of the knock down was quantified using the ImageJ software (NIH). The CK2 band intensity was normalized to the GAPDH band intensity and the knock down efficacy was determined relative to the mock condition.

### CK2 inhibition using CX-4945

CK2 was inactivated using the CK2-specific inhibitor CX-4945 (0308, Advanced Chemblocks). HeLa cells were seeded and transfected with CFP-VAP DNA the following day (for immunofluorescence assay only). 24 hours post-transfection, cells were infected with the *C. trachomatis incV* mutant expressing IncV-3xFLAG under the aTc inducible promoter at an MOI of 0.5-1. 18 hours post-infection, media containing 0, 0.625, or 10μM CX-4945 (final concentration) was added to each well and kept for the rest of the experiment. 2 hours after CX- 4945 addition (and 20 hours post-infection), media containing 20ng/mL aTc (final concentration) was added to induce IncV-3xFLAG expression. Cells were either collected in 2x Laemmli buffer and processed for western blot or fixed with 4% paraformaldehyde and processed for in immunofluorescence and confocal microscopy.

## Acknowledgments.

We thank Ted Hackstadt (NIH-Rocky Mountain Laboratories) and Mary Weber (University of Iowa) for the *incV::bla* strain; Claude Cochet and Odile Filhol-Cochet (Institut Albert Bonniot Département Réponse et Dynamique Cellulaires) for the YFP-CK2 constructs ; David Brautigan (University of Virginia) for the CX-4945 inhibitor.

We thank Hervé Agaisse and members of the Derré and Agaisse laboratories (University of Virginia) for providing feedback on the manuscript.

Funding sources were NIAID F31 AI136283 to RLM; Infectious Disease Training Grant T32 AI007046 to RLM, RJE, and SKD; NIAID R01 AI101441, NIAID R21 AI141841, and NIAID R01 AI162758 to ID.

## Supplementary Information

**Figure S1:**
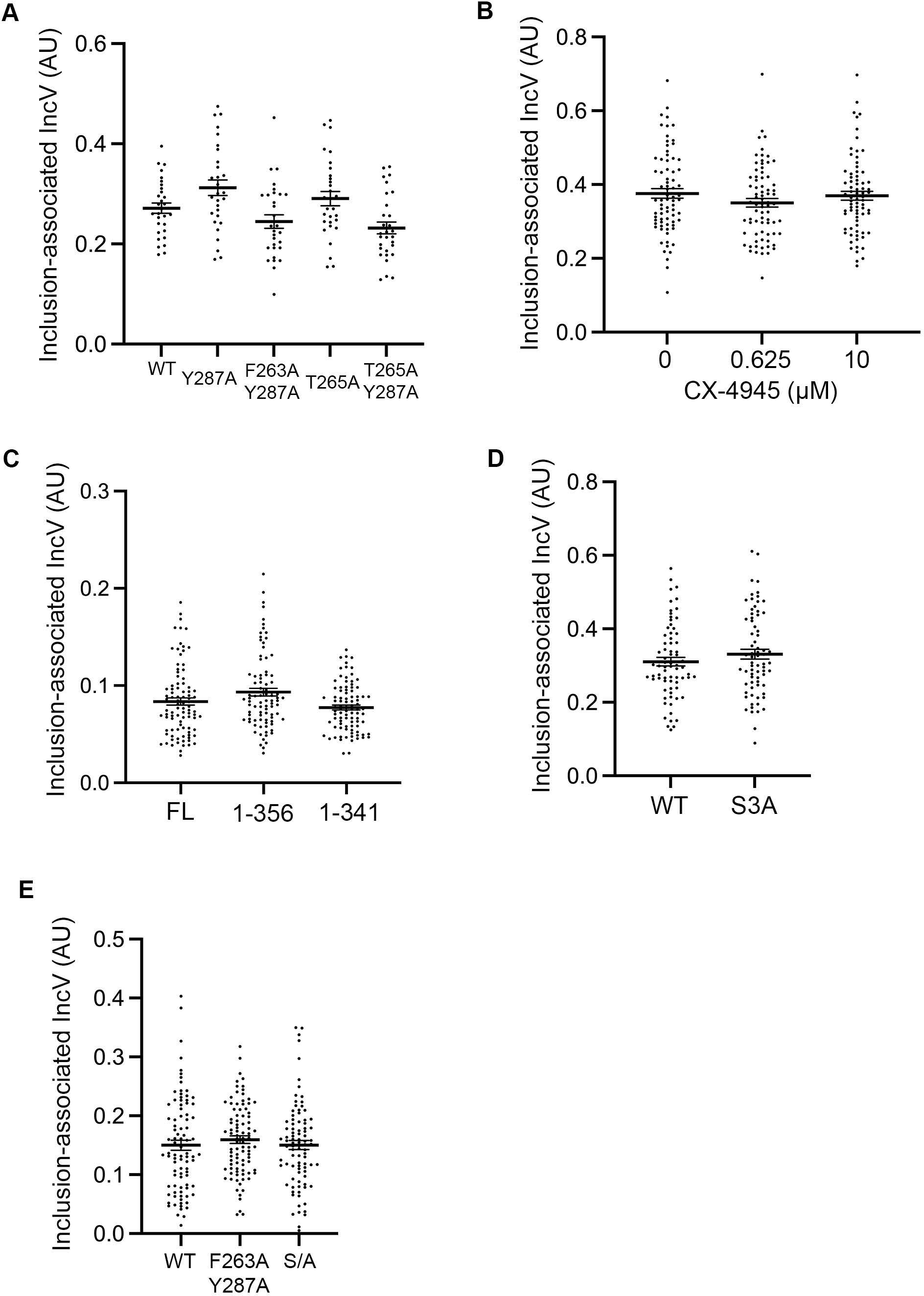
IncV inclusion localization is not affected upon CX-4945 treatment, truncation of IncV, or alanine substitution. (A-E) Quantification of the volume of the IncV-3xFLAG signal associated with the inclusion normalized to the volume of an object generated from the mCherry signal of the bacteria. Each dot represents one inclusion. The mean and SEM are shown. Student’s t-test (D) or One-way ANOVA and Tukey’s post hoc test (A-C, E) were performed comparing alanine substitution to wild type (A, D-E), drug treated cells to control cells (B), and truncations to full length (C).

**Figure S2:**
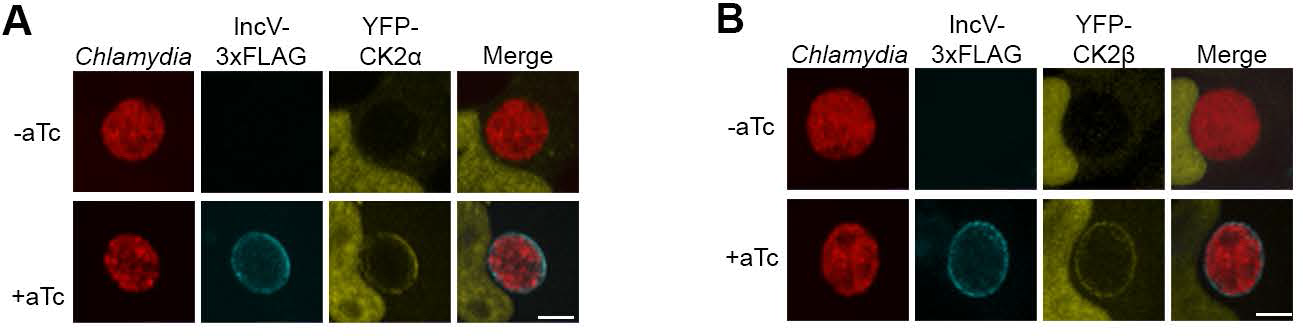
IncV recruits CK2 to the inclusion membrane. (A-B) 3-dimensional reconstruction of confocal images of HeLa cells overexpressing YFP-CK2α (A) or YFP-CK2β (B) (yellow) and infected with *C. trachomatis* expressing mCherry constitutively (red) and IncV-3xFLAG (blue) under the control of an anhydrotetracycline (aTc)-inducible promoter in the absence (-aTc) or presence (+aTc) of aTc. The merge is shown on the right. Scale bar is 5μm.

**Figure S3:**
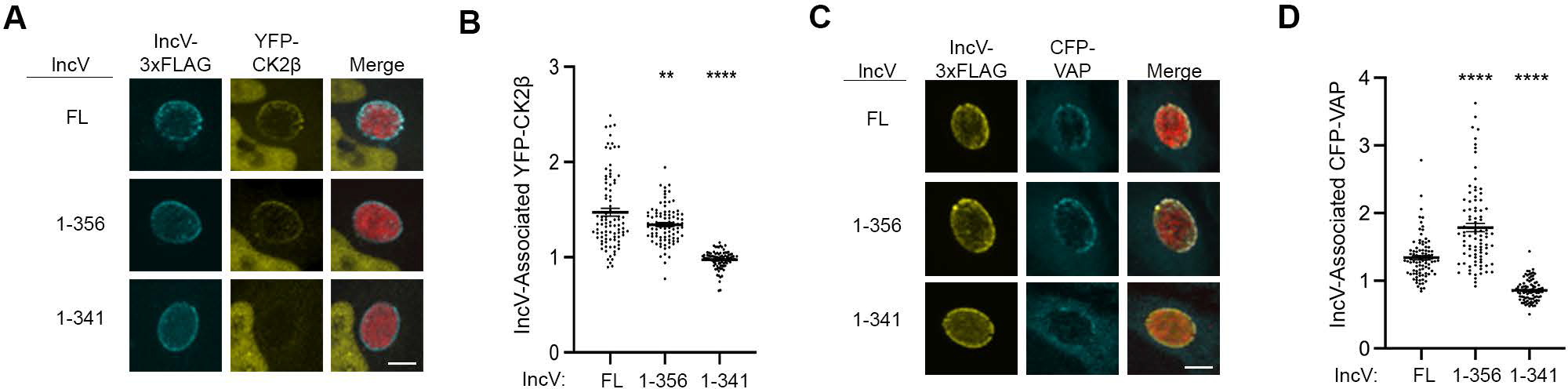
A C-terminal domain of IncV mediates VAP recruitment to the inclusion . (A) 3- dimensional reconstruction of confocal images of HeLa cells expressing YFP-CK2β (yellow) and infected with a *C. trachomatis incV* mutant expressing mCherry constitutively (red) and IncV-3xFLAG (full length (FL) or truncated (1-356, or 1-341) (blue) under the control of an aTc-inducible promoter in the presence of aTc. The merge is shown on the right. Scale bar is 5μm. (B) Quantification of the mean intensity of YFP-CK2β within an object generated from the IncV- 3xFLAG signal and normalized to the mean intensity of YFP-CK2β in the cytosol. Each dot represents one inclusion. Data show the mean and SEM of a combination of three independent experiments. One-way ANOVA and Tukey’s post hoc test was performed comparing truncations to full length. ** *P* <0.01, **** *P* <0.0001. (C) 3-dimensional reconstruction of confocal images of HeLa cells expressing CFP-VAP (blue) and infected with a *C. trachomatis incV* mutant expressing mCherry constitutively (red) and IncV-3xFLAG (full length (FL) or truncated (1-356, or 1-341) (yellow) under the control of an aTc-inducible promoter in the presence of aTc. The merge is shown on the right. Scale bar is 5μm. (D) Quantification of the mean intensity of CFP- VAP within an object generated from the IncV-3xFLAG signal and normalized to the mean intensity of CFP-VAP in the cytosol. Each dot represents one inclusion. Data show the mean and SEM of a combination of three independent experiments. One-way ANOVA and Tukey’s post hoc test was performed comparing truncations to full length. **** *P* <0.0001

**Figure S4:**
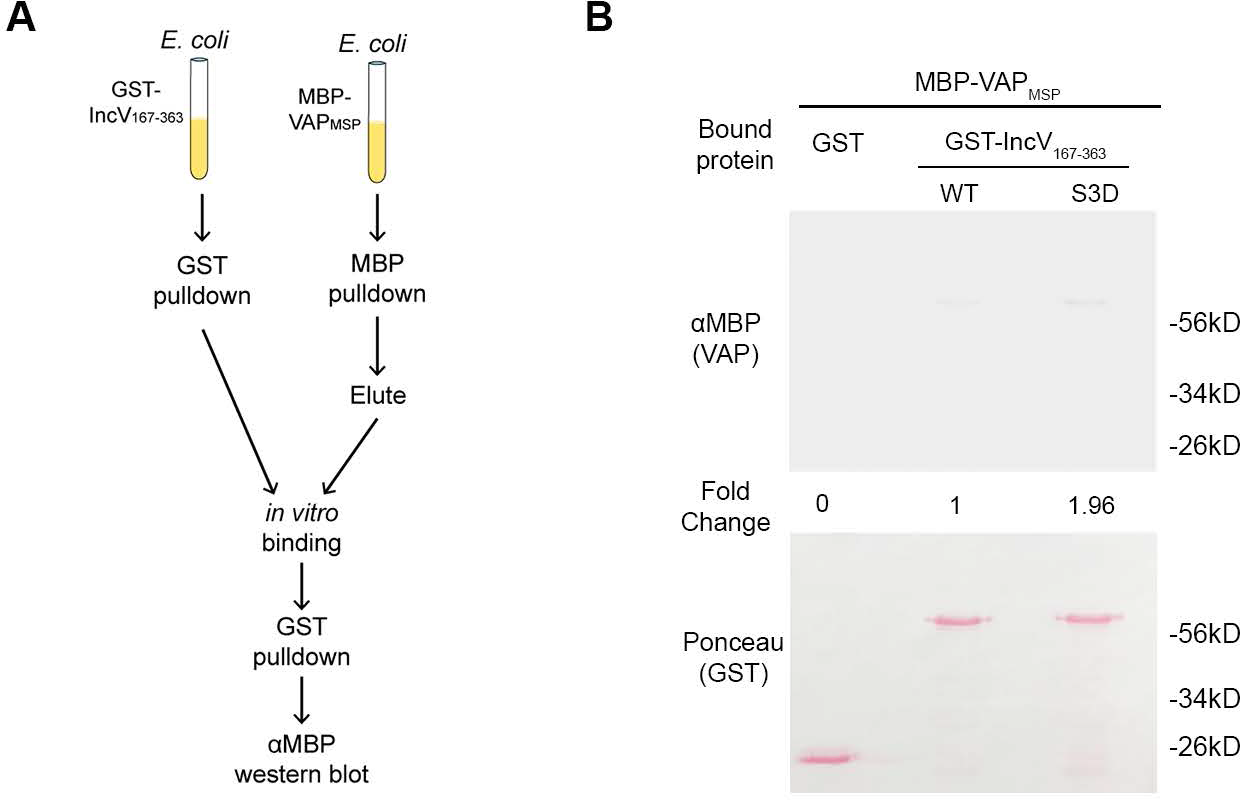
Phosphomimetic mutation of three serine residues in the C-terminal domain of IncV is not sufficient to promote the IncV-VAP interaction. (A) Schematic depicting the experimental setup for results in B. (B) *In vitro* binding assay using GST, GST-IncV_WT_, or GST- IncV_S3D_ purified from *E. coli*, immobilized on glutathione beads, and combined with MBP-VAP purified from *E. coli.* The top panel was probed with anti-MBP and the bottom panel was the same membrane stained with Ponceau S to detect the GST construct. Note that the IncV and VAP constructs, only include the cytosolic domain of IncV (aa 167-363) and the MSP domain of VAP, respectively.

**Figure S5:**
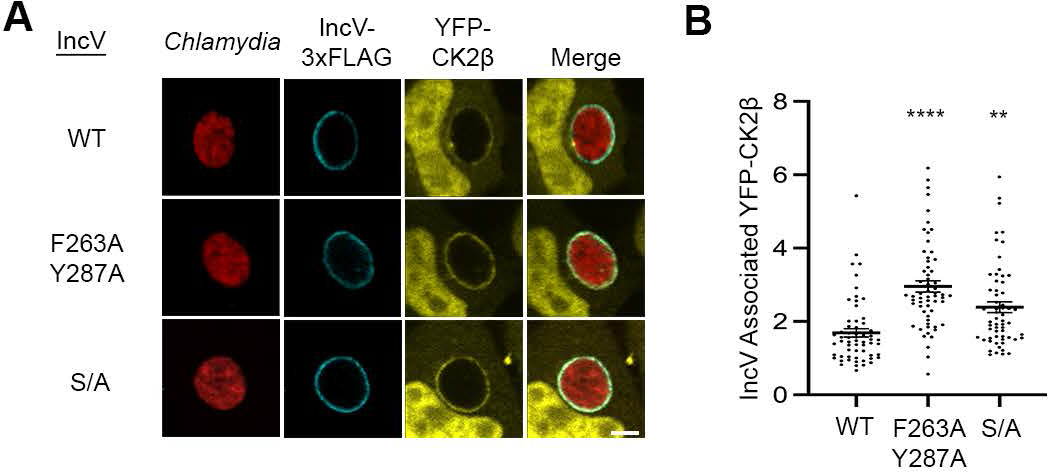
Alanine substitution of residues in position 2 of IncV FFAT motifs or of the serine rich tracts upstream of IncV FFAT motifs does not affect IncV-dependent CK2 recruitment to the inclusion. (A) Single plane confocal images of HeLa cells expressing YFP-CK2β (yellow) and infected with a *C. trachomatis incV* mutant expressing mCherry constitutively (red) and IncV_WT_-3xFLAG (WT), IncV_F263A/Y287A_-3xFLAG (F263A/Y287A), or IncV_S/A_-3xFLAG (S/A) (blue) under the control of an aTc inducible promoter. The merge is shown on the right. Scale bar is 5μm. (B) Quantification of the mean intensity of the YFP-CK2β within the IncV object normalized to the mean intensity of YFP-CK2β in the cytosol. Data show the mean and SEM of a combination of three independent experiments. One-way ANOVA with Tukey multiple comparisons test was performed to compare IncV_F263A/Y287A_ and IncV_S/A_ to IncV_WT_. ** *P* <0.01, \*\*\*\**P* < 0.0001.

**Figure 1 – source data 1:** Quantification of IncV-Associated YFP-VAP for Figure 1.

**Figure 2 – source data 1:** Uncropped, labelled blots for Figure 2A. Figure 2 **– source data 2:** Uncropped, labelled blots for Figure 2B. Figure 2 **– source data 3:** Uncropped, labelled blots for Figure 2D. Figure 2 **– source data 4:** Raw data for FLAG blot in Figure 2A. Figure 2 **– source data 5:** Raw data for FLAG blot in Figure 2B.

**Figure 2 – source data 6:** Raw data for Thiophosphate blot 1 in Figure 2D. Figure 2 **– source data 7:** Raw data for Thiophosphate blot 2 in Figure 2D. **Figure 2 – source data 8:** Raw data for Ponceau S blot 1 in Figure 2D. Figure 2 **– source data 9:** Raw data for Ponceau S blot 2 in Figure 2D.

**Figure 3 – source data 1:** Quantification of blot densities for Figure 3. Figure 3 – source data 2: Uncropped, labelled blots for Figure 3B. Figure 3 – source data 3: Uncropped, labelled blots for Figure 3D. Figure 3 – source data 4: Raw data for FLAG blot in Figure 3B. Figure 3 – source data 5: Raw data for Ponceau S blot in Figure 3B. Figure 3 – source data 6: Raw data for MBP blot in Figure 3D. Figure 3 – source data 7: Raw data for Ponceau S blot in Figure 3D.

**Figure 4 – source data 1:** Quantification of blot densities, line scan analysis and IncV-Associated CFP-VAP for Figure 4.

**Figure 4 – source data 2:** Uncropped, labelled blots for Figure 4A. Figure 4 **– source data 3:** Uncropped, labelled blots for Figure 4C. Figure 4 **– source data 4:** Raw data for FLAG blot in Figure 4A. Figure 4 **– source data 5:** Raw data for CK2 blot in Figure 4A. Figure 4 **– source data 6:** Raw data for GAPDH blot in Figure 4A. Figure 4 **– source data 7:** Raw data for FLAG blot in Figure 4C. Figure 4 **– source data 8:** Raw data for mCherry blot in Figure 4C. Figure 4 **– source data 9:** Raw data for Actin blot in Figure 4C.

**Figure 5 – source data 1:** Quantification of IncV-Associated YFP-CK2β and IncV-Associated CFP-VAP for Figure 5.

**Figure 5 – source data 2:** Uncropped, labelled blots for Figure 5F.

**Figure 5 – source data 3:** Raw data for FLAG and mCherry blots in Figure 5F.

**Figure 5 – source data 4:** Raw data for Actin blot in Figure 5F.

**Figure 6 – source data 1:** Quantification of IncV-Associated YFP-VAP and blot densities for Figure 6.

**Figure 6 – source data 2:** Uncropped, labelled blots for Figure 6D. Figure 6 **– source data 3:** Raw data for MBP blot in Figure 6D. Figure 6 **– source data 4:** Raw data for Ponceau S blot in Figure 6D.

**Figure S1 – source data 1:** Quantification of Inclusion-Associated IncV for Figure S1.

**Figure S3 – source data 1:** Quantification of IncV-Associated YFP-CK2β and IncV-Associated CFP-VAP for Figure S3.

**Figure S4 – source data 1:** Quantification of blot densities for Figure S4. **Figure S4** – source data 2: Uncropped, labelled blots for Figure S4B. **Figure S4** – source data 3: Raw data for MBP blot in Figure S4B. **Figure S4** – source data 4: Raw data for Ponceau S blot in Figure S4B.

**Figure S5 – source data 1:** Quantification of IncV-Associated YFP-CK2β for Figure S5.

## Notes

### Competing Interest Statement

The authors have declared no competing interest.

